# Predicted condensation-related signatures localize to two C-terminal regions of the prion protein across cervid PRNP variants associated with chronic wasting disease

**DOI:** 10.64898/2026.10.27.762346

**Authors:** Guy W. Dayhoff, Lyudmyla Dorosh, Holger Wille, Vladimir N. Uversky, Maria Stepanova

## Abstract

Genetic variation in cervid PRNP influences chronic wasting disease (CWD) susceptibility and progression, but it remains unclear whether disease-associated substitutions produce localized differences in sequence-derived properties relevant to prion protein (PrPC) condensation. We compared a white-tailed deer (WTD) reference sequence with 10 modeled variants on the same mature PrP scaffold (residues 24–233). For each sequence, we calculated intrinsic disorder, FuzDrop per-residue droplet-promoting propensity (pDP) and binding-mode entropy (Sbind), predicted residue-level pKa values, and a discrete-charge summary at pH 7.0. The overall profiles preserve a long N-terminal droplet-promoting region, whereas variant-associated differences are concentrated in selected intervals. In the FuzDrop outputs, residues 126–140 show allele-dependent changes in pDP near the droplet-promoting threshold, and residues 220–230 show variant-specific changes in Sbind, including lower Sbind for S225F/Y and higher Sbind for elk-like Q226E. These differences occur without large changes in overall intrinsic disorder. KaML pKa predictions identify near-neutral titratable sites that provide electrostatic context for these local predictor differences. Independent atomistic simulations of monomeric PrP further detected variant-sensitive backbone behavior and intramolecular contacts involving these intervals. Together, the analysis identifies residues 126–140 and 220–230 as computationally defined candidate intervals that overlap regions implicated independently in prion conversion or CWD fibril structure, motivating direct tests of cervid PrP condensation and maturation.

## Introduction

Prion diseases are fatal, transmissible neurodegenerative disorders caused by conformational conversion of the host-encoded prion protein (PrP^C^) into a misfolded, β-sheet-rich self-propagating isoform (PrP^Sc^).^1,2^ Chronic wasting disease (CWD) affects cervids, transmits within and between cervid species, and persists environmentally through shedding and adsorption to soils and plants.^1–5^ There is no approved vaccine or cure, and infection is invariably fatal.^1,2^

Host genetics, especially polymorphisms in *PRNP*, modify CWD susceptibility, incubation time, and disease progression.^2,6^ In white-tailed deer (WTD, *Odocoileus virginianus*), Q95H, G96S, and A116G have been linked to lower CWD case representation or delayed progression, while Q226K was associated with less advanced disease in a cohort study.^6–10^ In mule deer (*Odocoileus hemionus*), S225F is less frequently represented among CWD cases;^6,11^ in Rocky Mountain elk (*Cervus elaphus nelsoni*), M132L prolongs incubation relative to 132M;^12–14^ and in reindeer/caribou (*Rangifer tarandus*), studies of S138N and S225Y report differences in clinical disease or case representation, with sequence surveys providing additional context.^15–19^ Residue 226 also distinguishes deer-and elk-typical PrP contexts and influences CWD strain selection.^6,20^ These observations motivate a residue-resolved comparison of disease-associated cervid PrP variants.

Cervid PrP contains an intrinsically disordered N-terminal tail, including the octarepeat (OR) region and polybasic segments, and a structured C-terminal domain (Fig. 1).^21^ Sticker–spacer models offer a general framework for understanding interactions in low-complexity protein regions.^22,23^ More broadly, intrinsically disordered regions can support multivalent interactions relevant to phase separation.^24,25^ Experimental and review studies identify the PrP N terminus as a major contributor to PrP liquid–liquid phase separation (LLPS).^26,27^ Under the assay conditions used by Kamps et al., the OR and tryptophan residues contributed to phase separation, whereas histidines and copper were not required.^26^ Full-length recombinant PrP can also form droplets that mature toward Thioflavin T-positive, β-sheet-rich material under polymer–salt conditions.^28^ These studies establish N-terminal condensation as the biological background for the present analysis. Whether disease-associated substitutions in the C-terminal domain alter phase behavior remains unknown.

**Figure 1:**
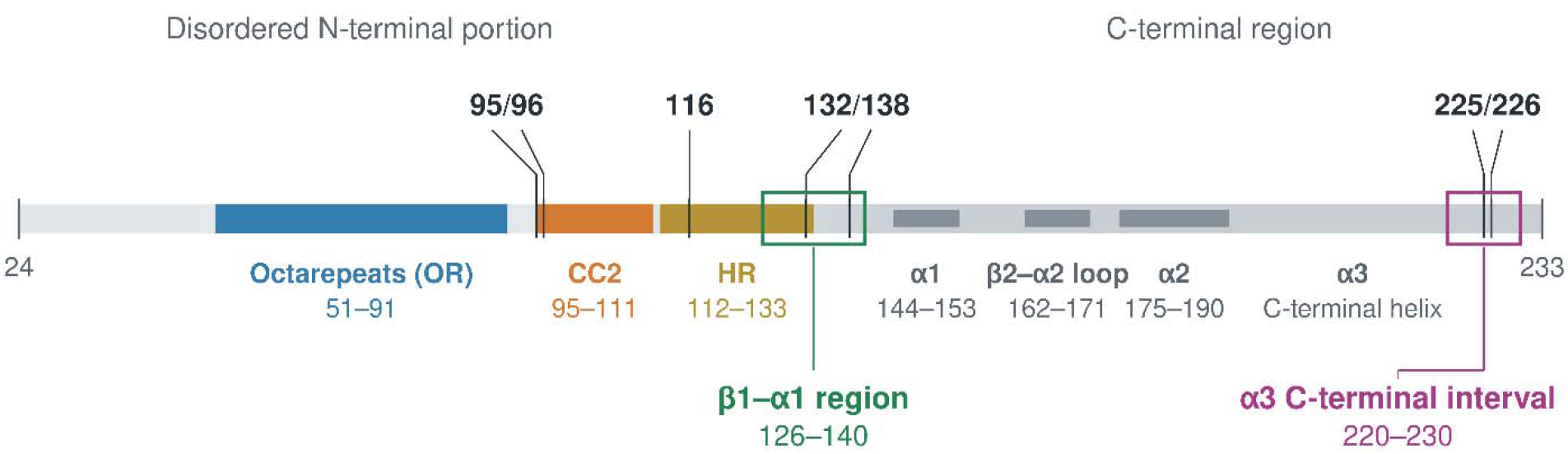
Cervid PrP sequence architecture and analyzed regions. Schematic of the white-tailed deer (WTD) PrP analytical sequence (residues 24–233), showing the octarepeat region (OR; 51– 91), charged cluster 2 (CC2; 95–111), hydrophobic region (HR; 112–133), selected secondary-structure elements, and modeled variant positions (95, 96, 116, 132, 138, 225, and 226). Green and magenta outlines highlight the two focal analytical intervals, residues 126–140 (β1–α1 region) and 220–230 (α3 C-terminal interval), respectively. All residue positions follow the common WTD PrP numbering frame.

Experiments comparing human Y145Stop PrP with full-length PrP provide additional context: under the conditions tested, the C-terminally truncated construct showed greater phase separation and matured into self-seeding amyloid.^29^

Within the PrP globular domain, the β2–α2 loop and surfaces spanning the α2–α3 loop and helix α3 contribute to templated conversion, species barriers, and strain selection. A single deer–elk difference at residue 226 shifts CWD strain propagation,^20^ and solution NMR structures of mule deer PrP show cervid-specific surface-charge features in this region.^21^ Cryo-EM structures of fibrils from naturally CWD-infected WTD further place the C-terminal lobe within the ordered PrP^Sc^ fibril core.^30^

Because the N-terminal contribution to PrP LLPS is established, the unresolved question is whether disease-associated sequence differences elsewhere in PrP produce localized predicted changes rather than broad shifts in global protein properties. A comparative study of cervid PrP variants found weak coupling among molecular-dynamics-derived backbone behavior, experimentally measured thermal stability, and spontaneous misfolding propensity, suggesting that a single global descriptor may not capture all variant-specific differences.^31^ Figure 1 summarizes the sequence architecture and analyzed regions.

Local electrostatics may contribute to variant-specific differences. Near-neutral titratable residues have side-chain pK_a_ values close enough to physiological pH that modest changes in pH or local environment can alter protonation state and charge, most notably for histidine. The N-terminal polybasic region is implicated in PrPSc binding and propagation, while experimental studies have examined metal-associated effects on PrP structure and condensation under defined conditions.^32–34^ Predicted pK values therefore provide residue-level electrostatic context for the analysis.

Here, we compare sequence-resolved computational profiles across a defined panel of cervid PRNP variants. RIDAO provides per-residue intrinsic-disorder estimates,^35^ FuzDrop provides p and S,^36^ and KaML provides predicted residue-level pK values.^37,38^ All profiles are mapped to the mature WTD PrP 24–233 sequence and compared with the WTD reference. We first examine the full sequence and then summarize predefined structural intervals. In these regional summaries, residues 126–140 were retained for their p_DP_ response, whereas residues 220–230 were retained as a separate C-terminal interval with variant-dependent S_bind_ differences and strong predicted pK_a_ variation. KaML predictions provide electrostatic context, and published LLPS, maturation, and structural studies provide biological context.^26,28^

## Materials and methods

### Sequence panel and variant provenance

The analysis included 11 PrP sequences: one white-tailed deer (WTD) wild-type (WT) reference, nine single-substitution variants, and one octarepeat deletion. Variants were selected from the cervid chronic wasting disease (CWD) literature to represent naturally occurring *PRNP* changes with published evidence of altered disease susceptibility, case representation, progression, incubation time, or strain selection. The sequence panel and the rationale for inclusion are summarized in Table 1.

**Table 1:** Cervid PrP sequence panel and analytical classification.

| Sequence | Literature provenance | Analytical role | Basis for inclusion |
|---|---|---|---|
| WT | White-tailed deer | Reference | WTD sequence used as the common scaffold and baseline for all comparisons.<sup>39</sup> |
| Q95H | White-tailed deer | Protective subset | Association with reduced case representation or delayed disease progression.<sup>7–10</sup> |
| G96S | White-tailed deer | Protective subset | Association with reduced case representation, altered tissue deposition, or prolonged disease course.7–10 |
| A116G | White-tailed deer | Protective subset | Association with reduced infection odds or less advanced disease in natural-host studies.<sup>6,10</sup> |
| M132L | Elk | Protective subset | Association with prolonged incubation and partial protection relative to 132M.13–15 |
| S138N | Reindeer/caribou | Protective subset | Experimental and field evidence of strong clinical protection; subclinical infection has been reported in gene-targeted models.<sup>15,16</sup> |
| S225F | Mule deer | Protective subset | Reduced case representation in natural mule deer populations.6,12 |
| S225Y | Reindeer/caribou | Protective subset | Under-representation among CWD cases relative to Ser225.<sup>17–19</sup> |
| Q226K | White-tailed deer | Protective subset | Lower disease stage among infected deer in one cohort; infection-odds trend was nonsignificant.<sup>6,10</sup> |
| Q226E | Elk-typical | Strain-context comparator | Deer–elk difference at residue 226 implicated in CWD strain selection and propagation.<sup>20</sup> |
| ORdel | Reindeer | Deletion comparator | Naturally occurring 24-base-pair octarepeat deletion associated with altered CWD risk.<sup>17</sup> |

All substitutions and the deletion were introduced into the common WTD scaffold described below. These classifications summarize the rationale for inclusion. The protective-subset label is descriptive and does not imply complete resistance or equivalent evidence across variants.

For the regional summaries, Q95H, G96S, A116G, M132L, S138N, S225F, S225Y, and Q226K were also analyzed as the eight-variant “protective subset,” a descriptive group defined by published evidence of reduced susceptibility or case representation, delayed progression, or prolonged incubation. The type and strength of evidence differ among variants; the term does not imply complete resistance. Q226E was analyzed separately as an elk-typical strain-context comparator, and ORdel as a deletion comparator. Each change was modeled individually on the same WTD reference scaffold, so species labels indicate the literature provenance of each substitution; all analyses use the common WTD scaffold. The panel was restricted to the variants and comparators summarized in Table 1.

### Study design and workflow

The analysis followed a fixed sequence. First, each substitution or deletion was introduced independently into the WTD reference sequence. Second, full-length per-residue profiles were generated for intrinsic disorder, FuzDrop condensation-related descriptors, and predicted pK_a_ values. Third, each variant profile was mapped to the WT coordinate system and expressed as a residue-wise variant–WT difference. Fourth, predefined sequence and structural intervals were summarized across the complete non-WT panel and the protective subset. Finally, electrostatic summaries and structural visualizations were used to interpret the sequence-level patterns. RIDAO, FuzDrop, and KaML-ESM provide the primary sequence-based outputs; KaML-CBTree provides supporting structure-informed pK_a_ context. Focal intervals were selected from the sequence-based regional summaries; structural models and published experiments were used during biological interpretation. Supplementary Fig. S3 summarizes the workflow.

### Sequence construction, numbering, and alignment

All analyses use mature cervid PrP numbering, spanning residues 24–233. In this frame, residue 24 is the cysteine that begins the mature N terminus (“CKKRPK. . .”) immediately after signal-peptide cleavage; residues 232–233 are the first two residues of the cleavable GPI-anchor signal. Unless otherwise noted, residue positions and axis labels refer to this 24–233 numbering scheme. The WTD WT sequence (NCBI QAU19527.1; UniProt Q863E9) served as the reference scaffold.^39^ Each substitution sequence differed from WT only at the stated position. ORdel was constructed by removing one canonical PHGGGWGQ repeat unit, corresponding to residues 56– 63 on the WT axis.

Cross-species polymorphic sites at positions 95, 96, 116, 132, 138, 225, and 226 align 1:1 across the cervid sequences considered here. Some PrP literature uses a human-centered convention in which the mature N terminus is labeled residue 23 and begins “KKRPK. . .”. All main-text analyses use the cervid 24–233 frame. Human residue numbers cited elsewhere retain the convention of their source studies and should not be interpreted using a fixed offset; local sequence correspondences are described in the Supplementary Methods.

All per-residue outputs were mapped to the common 24–233 axis. Substitution variants map directly to WT positions. ORdel was globally aligned to WT using the Needleman–Wunsch algorithm^40^ with a match score of +1, mismatch score of −1, and gap penalty of −2. Deleted positions were treated as missing: no values were interpolated across the deletion, and downstream residues were reported at their WT-aligned coordinates.

For metric *m*, the residue-wise difference for variant *v* at aligned WT position i was defined as

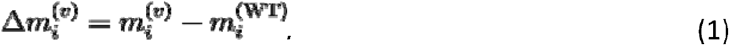

The term “variant–WT difference” is used throughout for this quantity.

### Prediction methods

Residue-level intrinsic disorder was predicted with Rapid Intrinsic Disorder Analysis Online (RIDAO), which integrates six per-residue disorder predictors: PONDR VLXT, PONDR VL3, PONDR VSL2B, PONDR-FIT, IUPred-short, and IUPred-long.^35^ Intrinsic disorder was summarized using the RIDAO mean disorder profile (MDP), the arithmetic mean of the six predictor outputs at each residue. MDP ranges from 0 to 1; MDP ≥ 0.50 was used as a descriptive disorder threshold.

Liquid–liquid phase-separation properties were predicted with FuzDrop.^36,41^ Four outputs were retained for each sequence:

- **Residue droplet-promoting propensity (p_DP_)**. p_DP_ ranges from 0 to 1 and estimates whether a residue promotes a droplet state within the FuzDrop framework. It combines the predicted probability of disorder in the free state with the predicted probability of disordered, or “fuzzy,” binding. Residues with p_DP_ ≥ 0.60 were classified as droplet-promoting.
- **Droplet-promoting regions (DPRs).** DPRs are contiguous segments of at least 10 residues with p_DP_ ≥ 0.60; FuzDrop additionally flags segments of at least 25 residues.
- **Binding-mode entropy (S_bind_, in nats).** S_bind_ describes predicted diversity of binding modes; larger values indicate greater context dependence. S_bind_ = 2.20 was used as a visual guideline for context-dependent regions (CDRs). CDR segments within a DPR were designated hotspots.
- **Protein-level LLPS probability (p_LLPS_**). p_LLPS_ is the sequence-level FuzDrop score in the range 0–1. Full per-residue p_DP_and S_bind_profiles, DPR/CDR/hotspot spans, and scalar p_LLPS_ values were retained. RIDAO MDP and FuzDrop outputs were analyzed separately.

p_LLPS_ was used as a comparative sequence-level model score across the sequence panel. Differences in p_LLPS_ were used to compare sequences and to motivate residue-level inspection of p_DP_ and S_bind_.

#### Additional phase-separation predictors

The same 11 mature PrP sequences were also evaluated with PScore,^42^ catGRANULES2,^43^ and DeePhase,^44^ which use feature sets distinct from FuzDrop. These predictors provided additional sequence-level comparisons across the same panel. Sequence-level scores and WT-referenced differences are reported in Supplementary Table S4, and residue-level PScore profiles are shown in Supplementary Fig. S4.

KaML-ESM provides sequence-only, residue-level pK_a_ predictions from protein-language-model embeddings. The model was trained using experimental pK_a_ shifts from PKAD-3, with separate acidic and basic channels and structure-informed pretraining from KaML-CBTree.^37,38^ We used the recommended ESM2 layer-31 embeddings and an ensemble of 200 multilayer perceptrons. Predictions were obtained for Asp, Glu, His, Cys, Tyr, and Lys; Arg pK_a_ is not modeled.

KaML-ESM values were analyzed as predicted residue-level pK_a_ values and variant–WT pK_a_ differences. They provided residue-level electrostatic annotations for the variant comparisons.

KaML-CBTree provides structure-informed pK_a_ predictions using features for solvent exposure, hydrogen-bond and salt-bridge opportunities, local structure or disorder, and electrostatic environment.^38^ A separate AlphaFold 3 single-chain structure was generated for each of the 11 PrP sequences and used as the corresponding KaML-CBTree input.^45^ The ORdel structure contained the same repeat deletion used in the sequence analyses. KaML-CBTree profiles provide the structure-informed pK_a_ analysis, while RIDAO, FuzDrop, and KaML-ESM were calculated independently.

### Derived electrostatic summaries

A conservative discrete-charge summary was calculated on the 24–233 axis at pH 7.0 using a centered seven-residue sliding window. For residue *i*, windowed net charge per residue (NCPR) was the arithmetic mean of assigned integer charges within the centered window; windows were truncated at sequence termini. Asp and Glu were assigned −1, Lys and Arg were assigned +1, and His was assigned +1 only when its KaML-ESM-predicted pKa exceeded 7.0; otherwise His was assigned 0. Tyr and Cys were treated as neutral. Charged-site counts were calculated from the same assignments.

We used this discrete-charge convention to limit propagation of predicted pK_a_ uncertainty.^37,38^ The charge summaries represent the apo state. Copper-dependent effects are discussed as potential modifiers of local protonation equilibria.^34^

### Regional summaries and selection of focal intervals

Regional summaries were calculated only after the full-length profiles had been generated. We compared the following predefined intervals on the 24–233 sequence: 24–50, 51–91, 92–120, 121–125, 126–140, 141–143, 144–153, 154–161, 162–171, 172–174, 175–190, 191–198, 199– 206, 207–219, 220–230, and 231–233. The overlapping 121–140 interval was also included to test whether the response extended from the pre-β1 linker through the β1–α1 region. Supplementary Tables S1–S2 give the structural labels used for these intervals.

For N-terminal comparisons, the OR was defined as residues 51–91. Residues 92–120 span charged cluster 2 (CC2; 95–111) and the N-terminal portion of the hydrophobic region (HR; 112–133); we refer to this explicitly defined interval as the CC2–HR neighborhood. The Supplementary Information also examines residues 86–91 and 92–97 to localize changes around the OR/CC2 transition.

For each metric *m*, variant *v*, and region *r*, the signed regional mean was calculated over aligned positions with valid values:

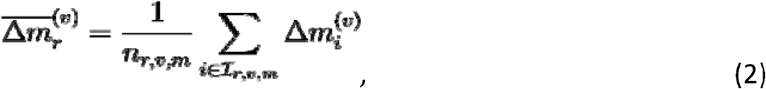

where I*r,v,m* is the set of aligned positions in region *r* with a value for metric *m*, and *n*r,v,m is the number of included positions. Signed means were used in regional heat maps and cross-metric plots. To compare the magnitude of regional responsiveness independently of direction, the mean absolute variant–WT difference was calculated as

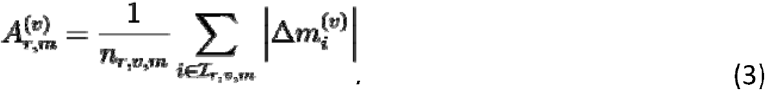

For *p*_DP_ and *S*_bind_, all aligned residues in the interval were included. For *pK_a_*, only aligned site with valid KaML predictions were included. Deleted ORdel positions were omitted. Region-level magnitudes were then summarized using the median across variants:

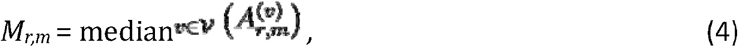

where V was either the complete set of 10 non-WT sequences or the protective subset. These metric-specific summaries are reported in Supplementary Tables S1–S2. Each metric wa evaluated separately during interval selection.

The focal intervals were chosen descriptively from the metric-specific regional summaries. When overlapping intervals captured the same local response, the narrower interval wa retained if it showed the larger median absolute response for the relevant metric. The selected intervals organize the focused analyses below, while Supplementary Tables S1–S2 retain the complete sequence-wide regional summaries. No classifier, significance test, or cross-metric composite score was used for interval selection.

#### Descriptive graphical summaries

Regional heat maps report the signed means from Eq. 2. For the per-variant p_DP_–S_bind_ summaries within regions 126–140 and 220–230, points represent the median residue-wise difference for each metric. Horizontal and vertical intervals are the 2.5^th^ and 97.5th percentiles of median values obtained from 10,000 bootstrap resamples of residues within the region. These intervals describe dispersion among residues, not confidence intervals based on independent biological replicates.

The cross-metric summary plot places mean Δp_DP_ for residues 126–140 on the x-axis and mean ΔS_bind_ for residues 220–230 on the y-axis for each non-WT sequence; WT is shown at the origin. Error bars are descriptive standard errors across residues within each interval. Marker shape encodes mean ΔMDP over residues 220–230; the ΔMDP = 0 label in Fig. 6 denotes a near-zero category (|ΔMDP| < 0.02). Spearman’s rank correlation coefficient was reported descriptively across the 10 non-WT sequences.

### Graphical summaries and structural visualization

For each sequence, the ordered pair (MDP*_i_*,p_DP,*i*_) was plotted for each residue *i*, and consecutive residues were connected in sequence order. These parametric maps provide a compact visualization of how intrinsic disorder and droplet-promoting propensity co-vary along PrP. Regional difference plots similarly compared Δp_DP_ with ΔMDP or ΔS_bind_ at aligned residues as descriptive visualizations.

A WTD WT AlphaFold 3 model spanning PrP residues 24–233 provided structural views of the sequence-derived results. RIDAO, FuzDrop, and KaML-ESM were calculated from sequence inputs; KaML-CBTree provided the structure-informed predictions.

For motif-centered *pK_a_* displays, the CBTree variant–WT *pK_a_* difference at each aligned ionizable site was averaged across the indicated variant group and displayed within named structural intervals. For structural coloring, the mean per-residue CBTree Δ*pK_a_* across the protective subset was mapped to aligned ionizable sites; quantitative summaries were limited to sites with comparable values.

Tyrosine correspondences link cervid positions to previously reported human PrP motifs; the quantitative analyses use cervid coordinates directly. The mapping criteria and complete residue correspondences are provided in the Supplementary Methods.

### Atomistic molecular dynamics of monomeric PrP

WT and nine single-substitution monomeric PrP systems (94–233) were each simulated in three independent 600-ns trajectories (18 μs total; GROMACS 2026.3/CHARMM36m).^46,47^ Regional backbone fluctuations, conformational populations, and solvent exposure were evaluated within residues 126–140 and 220–230; intramolecular contacts between these intervals were also measured. WT variation and mutation-excluded summaries provided controls (Supplementary Methods, Tables S5–S6, Fig. S5).

## Results

### Wild-type PrP defines the reference sequence landscape

Across the WTD reference sequence, FuzDrop identifies a long N-terminal droplet-promoting region that extends through much of the intrinsically disordered tail (Fig. 2b; Table 2). The structured C-terminal domain has lower p_DP_ overall, with local increases near residues 126–140 and at the C-terminal end of helix α3.

**Figure 2:**
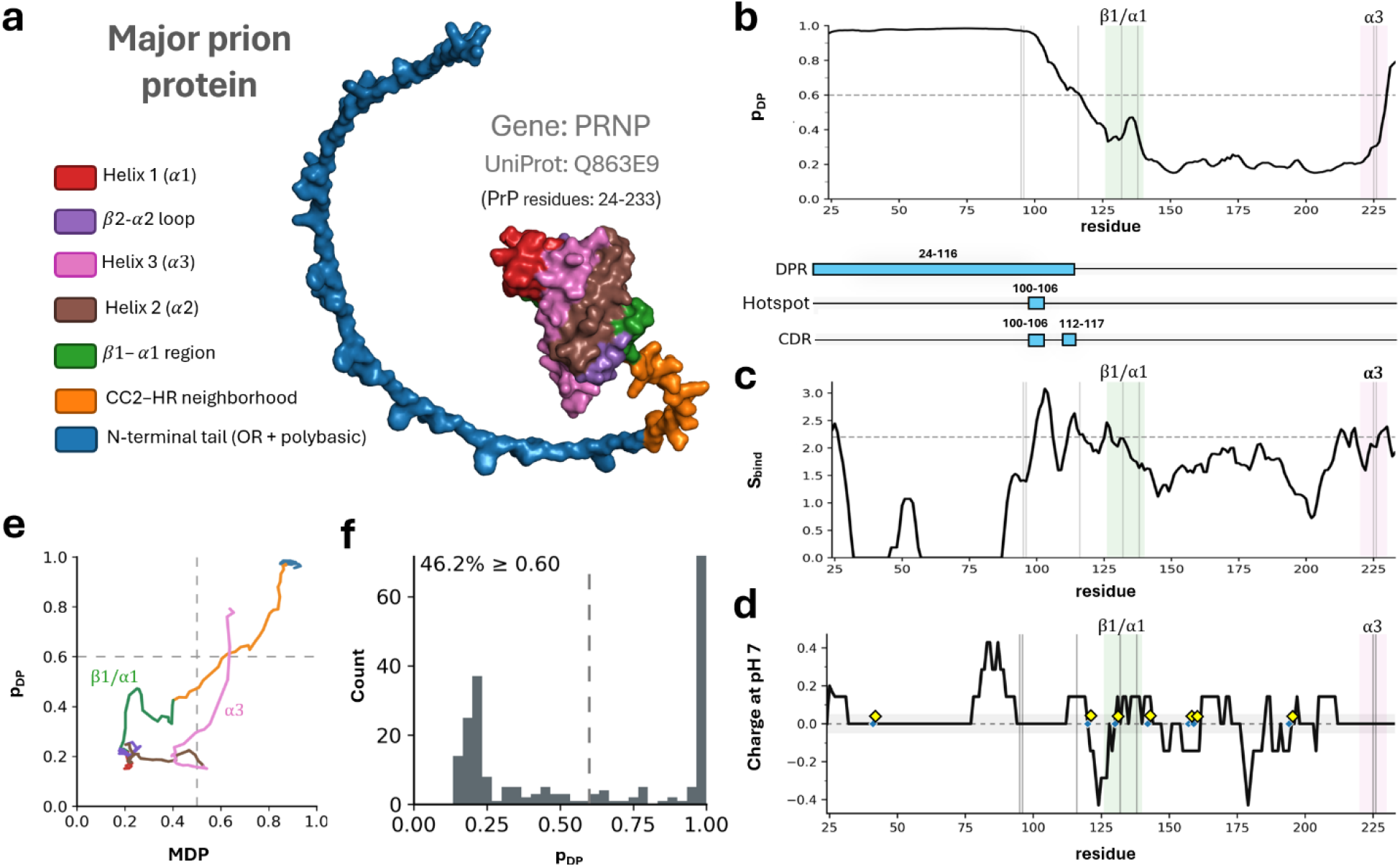
Wild-type cervid PrP defines the reference sequence landscape. **a**, AlphaFold 3 model of mature WTD PrP (24–233) shown for structural orientation. **b**, FuzDrop p_DP_ across residues 24–233; the 0.60 guideline and polymorphic sites 95, 96, 116, 132, 138, 225, and 226 are marked. Bars show droplet-promoting regions (DPRs), S_bind_ hotspots within DPRs, and context-dependent regions (CDRs; S_bind_ ≥ 2.20). **c**, FuzDrop S_bind_ with the 2.20 guideline. **d**, Discrete electrostatics at pH 7.0 (Asp/Glu = −1, Lys/Arg = +1, His = +1 only when KaML-ESM pK_a_ > 7.0; Tyr/Cys neutral); diamonds mark His residues with predicted pK_a_ > 7.0. **e**, Sequence-ordered p_DP_ versus RIDAO MDP. **f**, p_DP_ histogram with the 0.60 guideline.

**Table 2:**
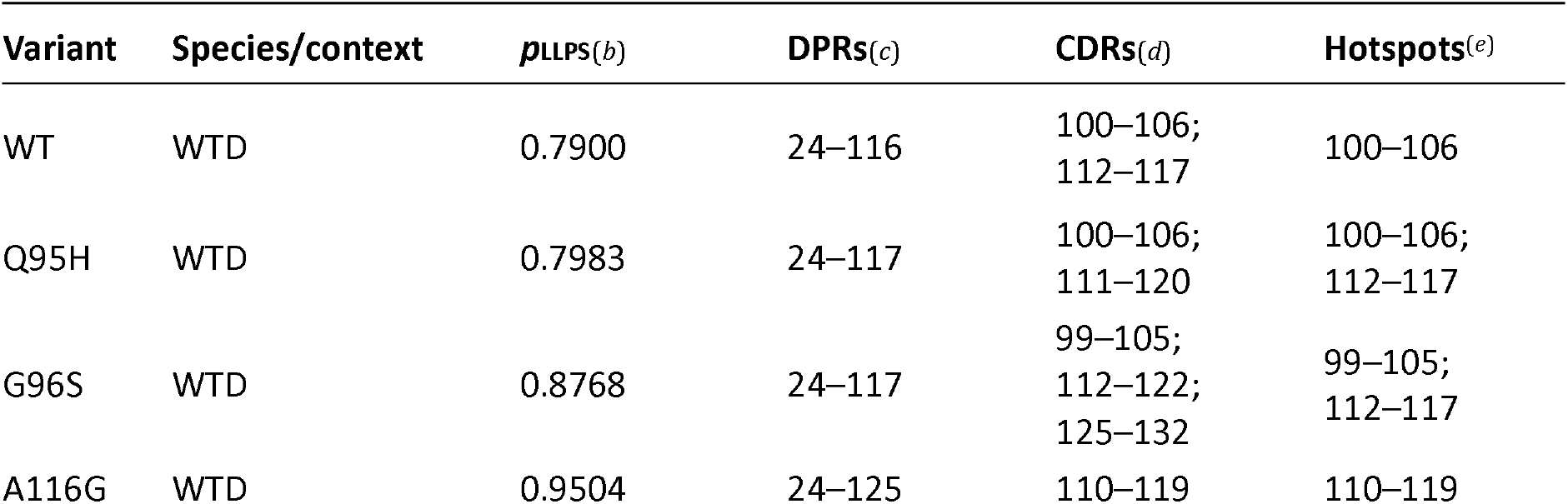
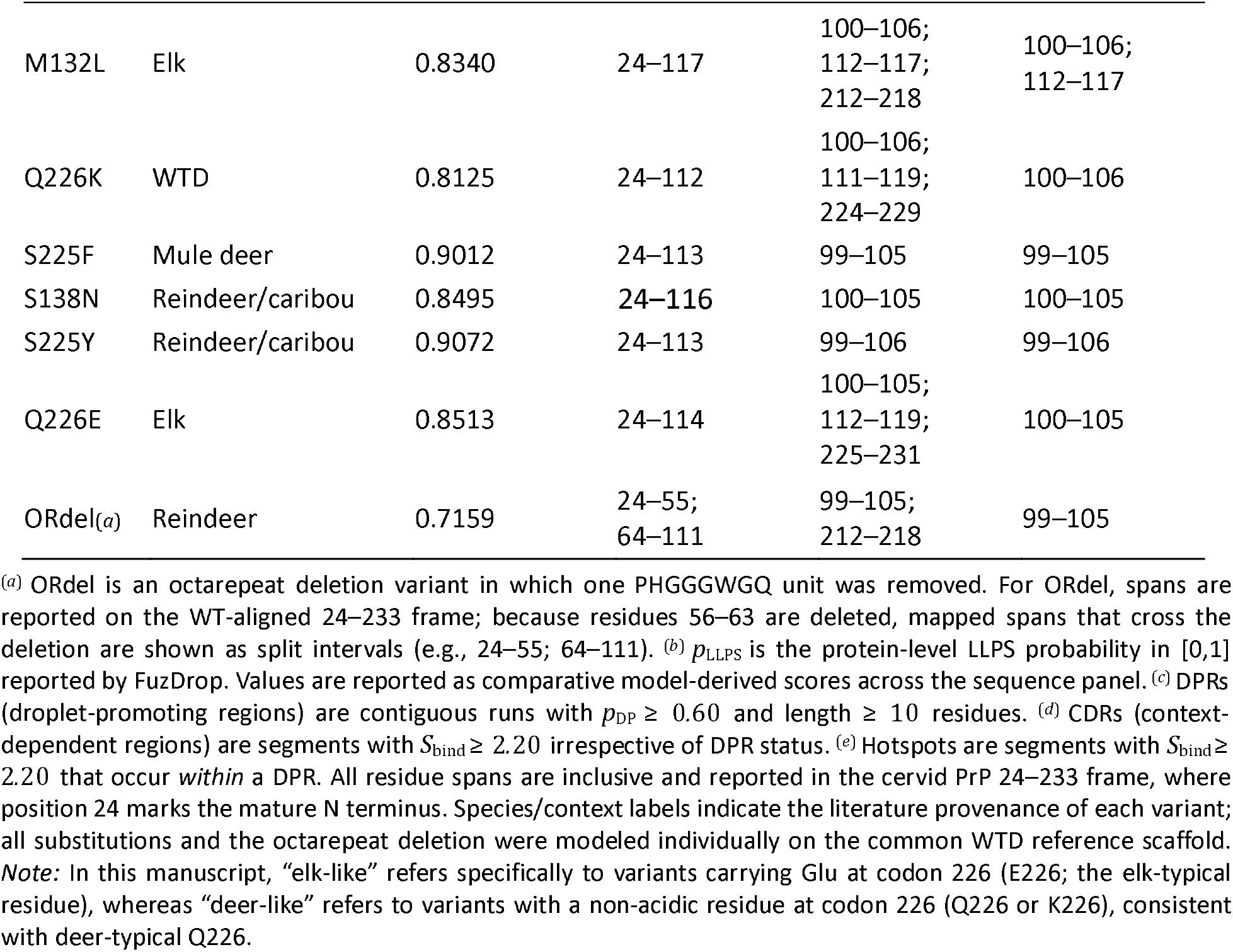
FuzDrop summary on the WTD scaffold (residues 24–233)

| **Variant** | **Species/context** | ***p*<sub>LLPS</sub>**<sub>(*b*)</sub> | **DPRs**<sub>(*c*)</sub> | **CDRs**<sub>(*d*)</sub> | **Hotspots**<sup>(*e*)</sup> |
|---|---|---|---|---|---|
| WT | WTD | 0.7900 | 24–116 | 100–106;<br>112–117 | 100–106 |
| Q95H | WTD | 0.7983 | 24–117 | 100–106;<br>111–120 | 100–106;<br>112–117 |
| G96S | WTD | 0.8768 | 24–117 | 99–105;<br>112–122;<br>125–132 | 99–105;<br>112–117 |
| A116G | WTD | 0.9504 | 24–125 | 110–119 | 110–119 |
| M132L | Elk | 0.8340 | 24–117 | 100–106;<br>112–117;<br>212–218 | 100–106;<br>112–117 |
| Q226K | WTD | 0.8125 | 24–112 | 100–106;<br>111–119;<br>224–229 | 100–106 |
| S225F | Mule deer | 0.9012 | 24–113 | 99–105 | 99–105 |
| S138N | Reindeer/caribou | 0.8495 | 24–116 | 100–105 | 100–105 |
| S225Y | Reindeer/caribou | 0.9072 | 24–113 | 99–106 | 99–106 |
| Q226E | Elk | 0.8513 | 24–114 | 100–105;<br>112–119;<br>225–231 | 100–105 |
| ORdel<sub>(*a*)</sub> | Reindeer | 0.7159 | 24–55;<br>64–111 | 99–105;<br>212–218 | 99–105 |

<sup>(*a*)</sup> ORdel is an octarepeat deletion variant in which one PHGGGWGQ unit was removed. For ORdel, spans are reported on the WT-aligned 24–233 frame; because residues 56–63 are deleted, mapped spans that cross the deletion are shown as split intervals (e.g., 24–55; 64–111). <sup>(*b*)</sup> *p*<sub>LLPS</sub> is the protein-level LLPS probability in [0,1] reported by FuzDrop. Values are reported as comparative model-derived scores across the sequence panel. <sup>(*c*)</sup> DPRs (droplet-promoting regions) are contiguous runs with *p*<sub>DP</sub> ≥ 0.60 and length ≥ 10 residues. <sup>(*d*)</sup> CDRs (context-dependent regions) are segments with *S*<sub>bind</sub> ≥ 2.20 irrespective of DPR status. <sup>(*e*)</sup> Hotspots are segments with *S*<sub>bind</sub> ≥ 2.20 that occur *within* a DPR. All residue spans are inclusive and reported in the cervid PrP 24–233 frame, where position 24 marks the mature N terminus. Species/context labels indicate the literature provenance of each variant; all substitutions and the octarepeat deletion were modeled individually on the common WTD reference scaffold.
*Note:* In this manuscript, “elk-like” refers specifically to variants carrying Glu at codon 226 (E226; the elk-typical residue), whereas “deer-like” refers to variants with a non-acidic residue at codon 226 (Q226 or K226), consistent with deer-typical Q226.

The sequence-wide regional summaries were then used to identify intervals with recurrent variant–WT differences (Supplementary Tables S1–S2). Residues 126–140 had the largest median absolute p_DP_ change in both the complete non-WT panel and the protective subset. The broader overlapping 121–140 interval was also responsive but had a smaller median absolute p_DP_ response, so residues 126–140 were retained for focused presentation. Residues 220–230 were retained as a separate C-terminal interval because they showed variant-dependent S_bind_ differences together with the largest median absolute predicted pK_a_ change in both variant sets. These intervals were selected from the full-length profiles.

The sequence-ordered p_DP_–MDP map provides a compact view of the same global organization (Fig. 2e). The N-terminal tail occupies the high-disorder, high-p_DP_ portion of the map, whereas the folded C-terminal core occupies a lower-disorder, lower-p_DP_ region. The p_DP_ profile approaches its threshold through residues 92–140, then rises near the C-terminal end of helix α3.

The remaining analyses therefore focus on variant–WT differences in p_DP_ across residues 126– 140 and in S_bind_ across residues 220–230, while retaining the full-sequence profiles for context. KaML pK_a_ predictions are used to annotate local electrostatics, and published structural and maturation studies are considered later in the Discussion.^20,28,30^

### Variant-resolved FuzDrop profiles show localized differences in residues 126– 140 and 220–230

Variant–WT differences are localized rather than uniform across PrP (Fig. 3). In residues 126– 140, several members of the protective subset, including G96S, A116G, and S138N, show higher p_DP_ across residues that lie near the FuzDrop droplet-promoting threshold (Fig. 3a,d). These changes occur without large shifts in RIDAO disorder propensity. In residues 220–230, S225F and S225Y show lower S_bind_, whereas elk-like Q226E shows higher S_bind_ (Fig. 3b,c,g). These S_bind_ differences indicate predicted changes in binding-mode diversity at the C-terminal end of helix α3. Protein-level p_LLPS_ scores differ among variants (Table 2), but do not identify the sequence positions of changes in p_DP_ or S_bind_. The regional profiles therefore retain positional information not represented by a single protein-level score.

**Figure 3:**
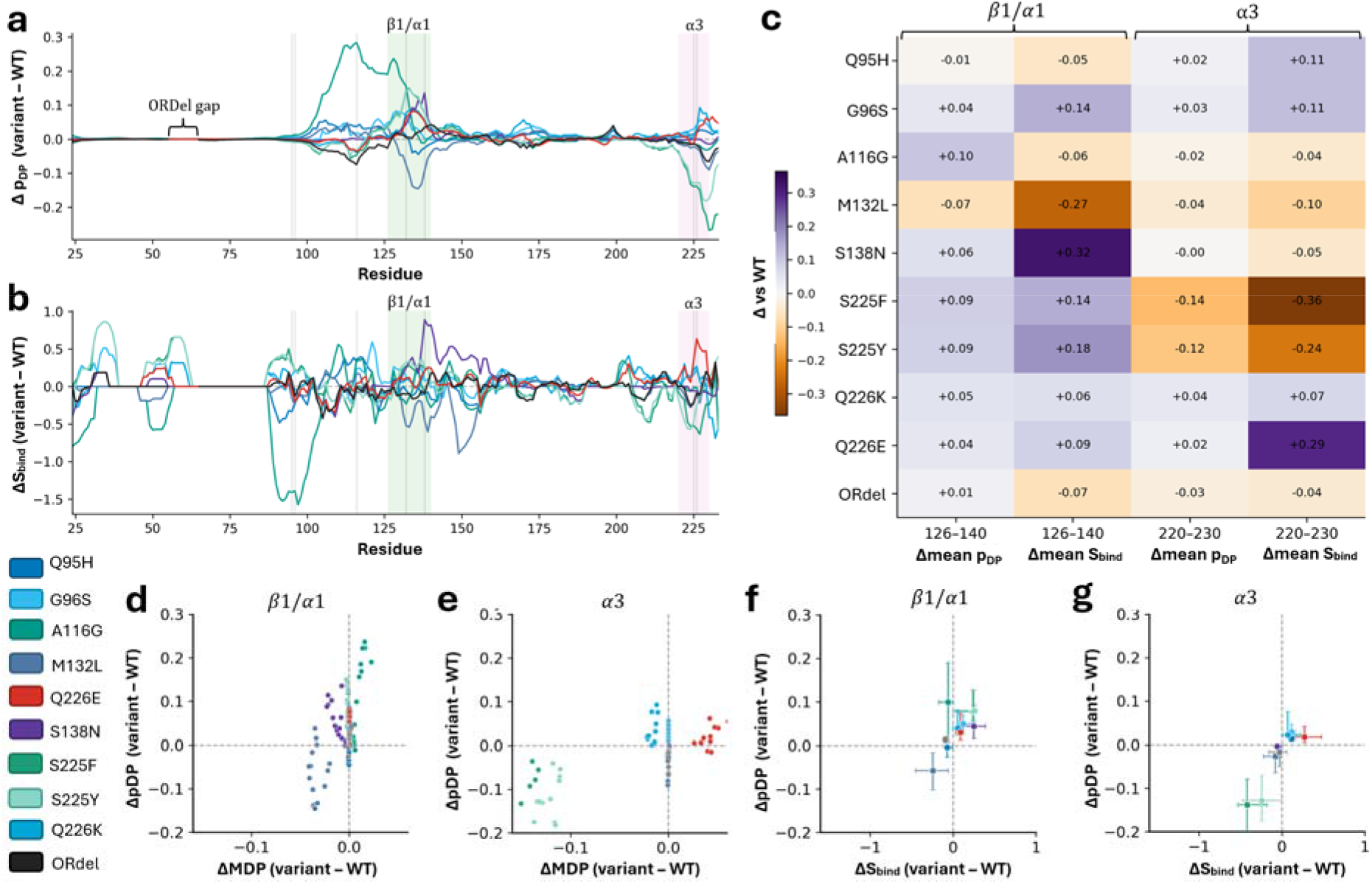
Variant-resolved FuzDrop differences are localized within residues 126–140 and 220– 230. **a**, Per-residue Δp_DP_ = p_DP_,variant − p_DP_,WT across residues 24–233. The zero line, focal intervals, and polymorphic sites are marked. **b**, Per-residue ΔS_bind_ = S_bind_,variant − S_bind_,WT. **c**, Region-averaged Δp_DP_ and ΔS_bind_ for residues 126–140 and 220–230. **d,e**, ΔMDP versus Δp_DP_ for residues 126–140 and 220–230, respectively. **f,g**, Per-variant median residue-wise Δp_DP_ and ΔS_bind_ within the two intervals. Horizontal and vertical intervals are the 2.5th and 97.5th percentiles from 10,000 bootstrap resamples of residues and summarize residue-level variation within each interval.

### Predicted pK_a_ values provide electrostatic context

KaML-ESM and KaML-CBTree provide sequence-only and structure-informed residue-level pK_a_ predictions, respectively (Fig. 4a). We use these values to identify titratable sites whose predicted protonation may be sensitive to local environment. Near-neutral sites occur within or near the intervals identified by the FuzDrop analysis, including the region around residue 226 (Fig. 4).^20,21^ The discrete pH 7.0 charge summary provides a simple sign-level view of the same electrostatic context (Fig. 2d). These pK_a_ predictions provide residue-level electrostatic annotations for the FuzDrop-defined intervals.

**Figure 4:**
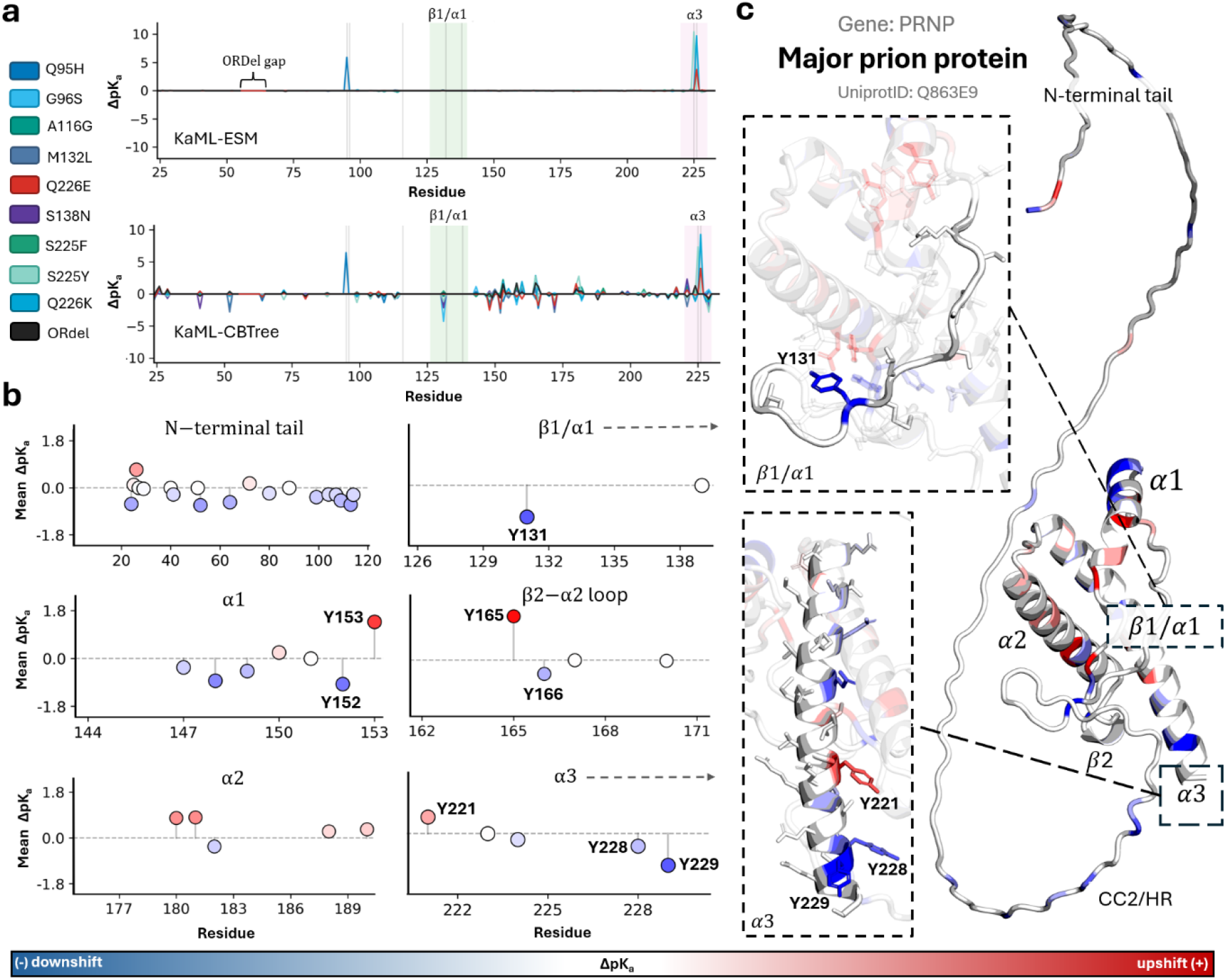
Predicted pK_a_ values provide residue-level electrostatic context. **a**, Per-residue ΔpK_a_ = pK_a_,variant − pK_a_,WT across residues 24–233 for KaML-ESM (sequence-only; top) and KaML-CBTree (structure-informed; bottom). The zero line, focal intervals, and polymorphic sites are marked. **b**, Mean KaML-CBTree ΔpK_a_ at aligned ionizable sites across the protective subset within the indicated sequence intervals. **c**, Mature WTD PrP colored by mean per-residue KaML-CBTree ΔpK_a_ across the protective subset. Labeled tyrosines correspond to sites discussed in the Supplementary Methods.

### Overall PrP organization is preserved across variants

The MDP, p_DP_, and S_bind_ profiles retain the same broad sequence organization across the variant panel: a highly disordered, droplet-promoting N-terminal tail followed by a lower-disorder C-terminal domain (Fig. 5a). The p_DP_–MDP trajectories are likewise similar at the global level (Fig. 5b). The main differences are local: p_DP_ varies through residues 92–140, whereas variants at positions 225 and 226 differ primarily in S_bind_ across residues 220–230 (Fig. 3).

**Figure 5:**
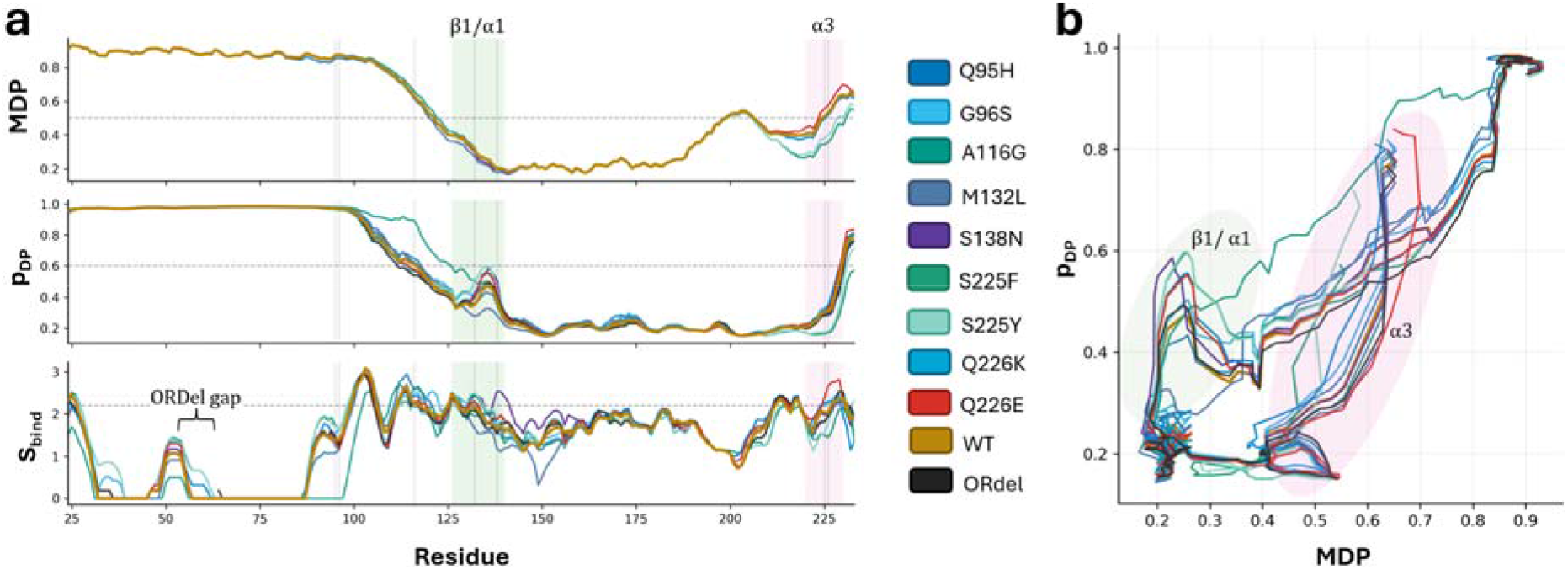
Overall PrP organization is preserved across variants. **a**, Per-residue RIDAO MDP (top), FuzDrop p_DP_ (middle), and FuzDrop S_bind_ (bottom) for all variants on the common 24–233 sequence. Guideline thresholds are MDP > 0.50, p_DP_ > 0.60, and S_bind_ > 2.20. The focal interval and polymorphic sites are marked. **b**, Sequence-ordered p_DP_ versus MDP for all variants. The global trajectory is similar across sequences, while local deviations are visible around residue 92–140 and 220–230.

### N-terminal and C-terminal intervals show distinct variant patterns

Q95H and G96S show their largest predicted differences near the OR/CC2 transition and within the CC2–HR neighborhood (residues 92–120). Across the full panel, the OR itself (51–91) changes little (median |Δp_DP_| = 0.0005; median |ΔS_bind_| = 0.0622), whereas residues 92–120 show larger differences (median |Δp_DP_| = 0.0167; median |ΔS_bind_| = 0.1465; Supplementary Table S1). The same ordering is present in the protective subset (eight variants; Supplementary Table S2).

For Q95H and G96S, the strongest residue-level changes occur around the OR/CC2 transition (Supplementary Figs. S1–S2). Both variants extend predicted context-dependent binding segments within residues 92–120, and G96S introduces an additional downstream CDR at residues 125–132. Their protein-level FuzDrop scores are also higher than WT (WT, 0.7900; Q95H, 0.7983; G96S, 0.8768; Table 2). These observations describe a distinct N-terminal predictor pattern rather than an α3-centered pattern.

Figure 6 summarizes the two C-terminal readouts across variants. Several members of the protective subset, including S225F and S225Y, combine higher mean p_DP_ in residues 126–140 with lower mean S_bind_ in residues 220–230. A116G and Q226K shift in the same direction but remain closer to the axes, whereas Q226E shows higher S_bind_ at the α3 C-terminus. Q95H and G96S show smaller changes along the α3 axis, consistent with the N-terminal pattern described above.

**Figure 6:**
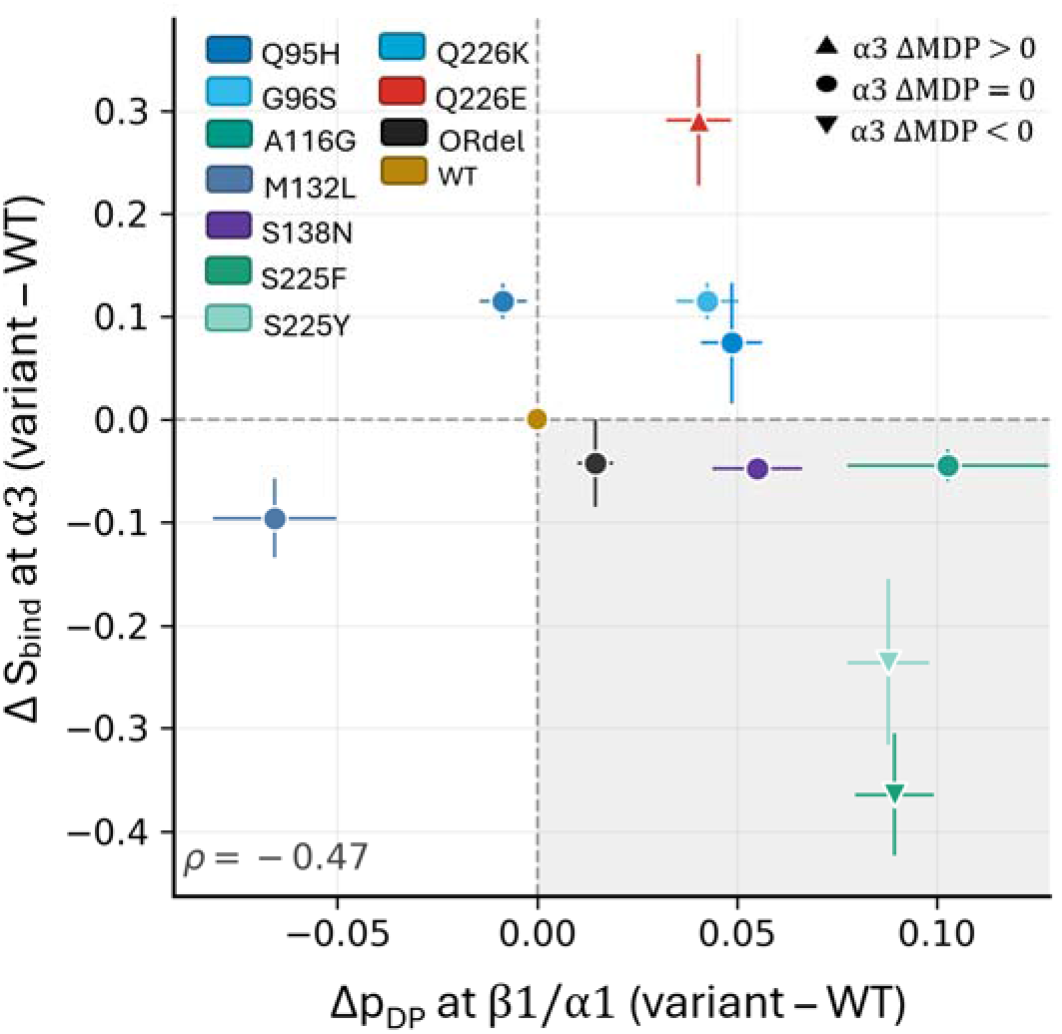
Cross-metric summary of variant differences in residues 126–140 and 220–230. Each point represents one variant. The x-axis is mean Δp_DP_ in residues 126–140 and the y-axis is mean ΔS_bind_ in residues 220–230, both relative to WTD WT. Error bars show descriptive standard errors across residues within each interval. Marker shape encodes mean ΔMDP over residue 220–230: upward triangle, increase; circle, approximately zero (|ΔMDP| < 0.02); downward triangle, decrease. The shaded lower-right quadrant marks Δp_DP_ > 0 together with ΔS_bind_ < 0. Across the 10 non-WT sequences, the descriptive Spearman correlation between the two axes i ρ = −0.47.

Residue-level comparisons also show that changes in intrinsic disorder are small and localized (Fig. 3d,e).

The β2–α2 and α2–α3 loop intervals were included in the same sequence-wide regional analysis even though none of the modeled substitutions lies within those loops. Both show smaller median variant–WT differences in p_DP_ and S_bind_ than residues 126–140 and 220–230 in the full panel and the protective subset (Supplementary Tables S1–S2). Their established roles in prion conversion therefore remain biologically important, but they are less responsive in the condensation-related descriptors examined here.

### Additional predictors show different sensitivities across the variant panel

The additional sequence-level phase-separation predictors do not reproduce the FuzDrop p_LLPS_ ordering across the 11 sequences (Supplementary Table S4). PScore shows its largest change for ORdel, catGRANULES2 varies modestly across the panel, and DeePhase values are narrowly distributed. Residue-level PScore profiles are shown in Supplementary Fig. S4. These methods use different feature sets and output definitions, and each predictor is interpreted according to its own score definition.

A separate published study of cervid PrP variants also found weak coupling among thermal stability, molecular-dynamics-derived behavior, and spontaneous misfolding propensity.^31^ The overlapping variants are compared in Supplementary Table S3. The M132L comparison is qualitative because Díaz-Domínguez et al. studied M132L in a Q226E background, whereas the present analysis models M132L on the WTD Q226 scaffold. Together, these comparisons show that the methods rank variants differently. The focal intervals therefore remain hypotheses derived from the residue-level FuzDrop analysis.

Q226E provides a concrete example of this divergence: it shows higher predicted Sbind in residues 220–230 here, while its published spontaneous-misfolding score does not increase relative to the Q226 reference.^31^

### Independent MD assessment of the focal intervals

Across all 30 monomeric trajectories, the structured core remained stable (mean core-backbone RMSD, 0.085–0.126 nm). The predefined intervals showed variant-dependent differences in backbone fluctuations, secondary-structure and backbone-dihedral distributions, and solvent exposure (Supplementary Tables S5–S6; Fig. S5). Mean RMSF in residues 126–140 was higher for G96S (0.179 nm) and S225Y (0.155 nm) than for WT (0.124 nm). In residues 220–230, mean RMSF was higher for S225F (0.182 nm) than for WT (0.148 nm). The corresponding ranges across the three trajectories overlapped WT.

Sequence variation also affected the intramolecular interface between residues 126–140 and 220–230. For S225F, hydrogen-bonded residue pairs averaged 0.474–1.443 per frame across the three trajectories (WT, 0.130–0.318), and heavy-atom contacting residue pairs averaged 0.620–1.581 per frame (WT, 0.325–0.465). A116G also showed increased physical-contact occupancy in each trajectory (0.549–0.734 per frame). The interface effects for S225F did not depend on contacts involving residue 225. Other variant effects were less consistent relative to WT variation; notably, the S138N R1 response and the Q226E R2 solvent-accessibility response varied across trajectories. Together, these observations demonstrate variant-sensitive monomeric conformational behavior and intramolecular interactions involving the sequence-prioritized intervals.

## Discussion

### Established N-terminal LLPS provides the biological context

The long N-terminal FuzDrop DPR is consistent with direct experiments showing that the PrP N terminus, including the octarepeats and tryptophans, is a major driver of droplet formation,^26^ and that full-length PrP droplets can mature toward β-sheet-rich material under polymer–salt conditions.^28^ Heterotypic phase separation has also been reported to suppress amyloid formation by the Y145Stop PrP variant, showing that condensate context can alter downstream aggregation.^48^ The N-terminal polybasic segment also contributes to PrPSc binding and efficient prion propagation.^32^ These observations provide biological context for the N-terminal predictions and make the N-terminal region a practical reference for interpreting local variant differences.

Under the polymer–salt conditions of Tange et al., aged full-length PrP gels acquired β-sheet-rich, sarkosyl-insoluble, and proteinase K-resistant features but lacked detectable seeding activity, emphasizing that condensate maturation is not equivalent to templating-competent prion material.^28^

### Experimental context for pH and metal effects

Non-octarepeat Cu(II) coordination in human PrP varies with pH and sequence context,^33^ and metal-dependent PrP condensation has been reported under some near-neutral buffered conditions.^34,49^ By contrast, Tange et al. observed amorphous aggregation rather than liquid droplets after CuSO addition under their polymer–salt conditions.^28^ Together, these differing observations emphasize that metal-associated PrP phase behavior must be interpreted in the context of the experimental conditions. Accordingly, the near-neutral predicted pK_a_ sites in residues 126–140 and 220–230 provide candidates for direct tests of pH- and metal-dependent effects. Changes in protonation of suitably titratable sites could redistribute local charge and modify interaction preferences without a substantial change in overall disorder.

### Independent evidence supports the biological relevance of the focal intervals

The sequence neighborhood containing cervid residues 126–140 has independent links to prion amyloid formation and conversion. In the human Y145Stop model, truncation experiments implicated residues around 138–141 (human numbering) in amyloid formation.^50^ The same truncated protein undergoes liquid–liquid phase separation and matures into self-templating amyloid,^29^ while solid-state NMR studies of human PrP23–144 amyloid identified an ordered core spanning approximately human residues 112–141 and showed that variation at residue 139 can alter fibril structure.^51^ A protective human G127V substitution within the corresponding region also produces localized structural and dynamic changes around the first β-strand while retaining the overall PrP fold.^52^ In cervids, S138N falls within the focal interval and prevented clinical CWD in gene-targeted mouse experiments while reducing measured PrP conversion; subclinical infection and shedding could nevertheless occur.^16^ Together, the human structural studies and cervid S138N data provide independent biological support for examining residues 126–140. Their specific involvement in full-length cervid PrP condensation remains to be tested. Residues 220–230 have more direct cervid-specific support. Natural WTD CWD fibrils contain a C-terminal arch spanning residues 215–229 that is stabilized in part by a Q220–Y229 hydrogen bond.^30^ The deer–elk Q226/E226 difference affects disease kinetics, PrP^Sc^ conformation, and strain selection in otherwise syngeneic gene-targeted mice.^20^ In recombinant cervid RT-QuIC, the α3-associated 225F, 225Y, and Q226K substrates show reduced amplification rates across multiple CWD isolates and seed dilutions.^53^ Díaz-Domínguez et al. likewise reported lower spontaneous-misfolding scores for S225F/Y than for Q226Q/E, although the global biophysical readouts in that study do not share a single variant ordering.^31^ A recent independent structure-based EmCAST analysis predicted S225F to be approximately neutral in folded mule deer PrP but incompatible with the natural CWD fibril conformation used in that analysis.^54^ Together, these studies provide independent structural and conversion-related evidence for the biological importance of residues 220–230. The specific relationship between the predicted Sbind differences and PrP condensation or condensate maturation remains an experimental question. The independent monomeric MD analysis complements this biological evidence. The simulations showed variant-dependent conformational and fluctuation differences involving residues 126–140 and 220–230. S225F and A116G also showed greater intramolecular contact occupancy between these intervals than the observed WT range in all three trajectories. These measurements do not directly assess PrP condensation or intermolecular association. Together with the established N-terminal role in PrP condensation, the findings identify C-terminal regions for subsequent experimental tests of assembly and conversion.

### Testable predictions

The sequence-resolved analysis yields three experimentally testable predictions.

i. Constructs spanning the CC2–HR neighborhood (residues 92–120) should show larger Q95H/G96S-dependent changes in condensation-related behavior than constructs restricted to the OR alone, with the largest predicted differences near residues 92–97 (Supplementary Fig. S2).
ii. Variants with larger predicted changes across residues 126–140 are expected to differ in the coupling between droplet formation and subsequent structural conversion under matched conditions. This can be tested by measuring phase separation, droplet aging, ThT kinetics, proteinase K resistance, and seeding as separate endpoints, with pH titration used to test the contribution of nearby titratable sites.
iii. Published RT-QuIC measurements already show reduced amplification rates for the α3-associated 225F, 225Y, and Q226K substrates.^53^ We therefore predict that S225F and S225Y will alter the transition from condensed PrP to conversion-competent aggregates relative to WTD WT or Q226E, rather than uniformly slowing every stage of amyloid growth. This can be tested by separating LLPS onset and partitioning, droplet-aging and FRAP measurements, nucleation lag, post-threshold ThT growth, insoluble fractions, proteinase K resistance, and seeding readouts.

These experiments should use matched phase-separation conditions across variants so that differences in initial droplet formation can be distinguished from differences in maturation. Additional tyrosine-based reporter hypotheses are provided in the Supplementary Information.

### Limitations

Each substitution or deletion was evaluated on the same WTD WT scaffold. This design isolates the computational response to a single sequence change, but it does not reproduce complete native-species haplotypes or epistatic interactions. Results for elk-, mule deer-, and reindeer-associated substitutions therefore represent single-variant effects in a WTD sequence background.

FuzDrop pLLPS, pDP, and Sbind are comparative computational descriptors. They do not directly estimate saturation concentration, condensate material state, maturation kinetics, infectivity, or CWD risk. KaML pKa values are also model-derived and are used here as electrostatic context. PScore, catGRANULES2, and DeePhase show different sequence-level sensitivities from FuzDrop, reflecting their different feature sets and output definitions. Independent structural and conversion studies support the biological relevance of the focal intervals, but they do not directly test the FuzDrop-derived condensation signatures in these modeled cervid variants. The two focal intervals were selected descriptively from sequence-wide regional summaries and are used to organize hypothesis generation. Other PrP regions may also contribute to condensation, maturation, or prion conversion. Available cervid evidence supports partial protection or delayed disease rather than complete resistance.^17^ The disease-associated variant group includes different forms and strengths of evidence. Cross-species tyrosine correspondences follow the supported local motif mappings described in the Supplementary Methods.^55^

The MD simulations sampled monomeric PrP residues 94–233, rather than condensed assemblies; their structural measurements do not determine LLPS propensity. Some regional differences were sensitive to trajectory sampling, and the 600-ns observation window does not establish complete equilibrium convergence.

## Conclusions

Across the WTD reference and 10 modeled variants, the overall disorder and condensation-related profiles were broadly preserved, but localized differences emerged in FuzDrop predictions. Residues 126–140 show allele-dependent p_DP_ differences near the droplet-promoting threshold, whereas residues 220–230 show variant-specific S_bind_ differences, including lower S_bind_ for S225F/Y and higher S_bind_ for elk-like Q226E. Predicted pK_a_ values identify near-neutral titratable sites that provide electrostatic context for both intervals.

These results identify specific residue-level hypotheses for experimental testing. Protein-level FuzDrop and the additional sequence-level phase-separation predictors capture different patterns from the residue-level FuzDrop analysis. Independently of FuzDrop, studies of the broader sequence neighborhood of residues 126–140 implicate it in amyloid structure and conversion, and the cervid S138N variant within this interval has demonstrated effects in gene-targeted models;^16,29,50–52^ residues 220–230 overlap a C-terminal arch in natural WTD CWD fibrils, while studies of variants at 225 and 226 have reported differences in seeded amplification or CWD strain behavior.^20,30,53^ Together, independent structural and conversion studies support prioritizing both intervals, while their specific roles in PrP condensation remain to be tested experimentally. Atomistic simulations also showed variant-dependent monomeric conformational behavior involving the prioritized intervals, including reproducible changes in their intramolecular interface for selected variants.

The analysis therefore prioritizes specific sequence intervals and reporter residues for direct testing with pH-controlled LLPS assays, droplet-maturation measurements, and aggregation or seeding readouts.^26,28^ PrP droplets and PrP^Sc^ are distinct states, while infectious CWD fibril structures provide structural context for the predicted condensation-related differences.^30^ The resulting framework is a reproducible starting point for testing how cervid sequence variation may alter local PrP interaction behavior. Candidate reporter sites include Y152/Y153 in helix α1 and Y228/Y229 near the C-terminal end of helix α3 (mature cervid numbering; Supplementary Methods).

## Data availability

All source data underlying the sequence-based figures and analyses are deposited on Zenodo and are publicly available. The permanent record is: https://doi.org/10.5281/zenodo.17253991. The archive includes:

- **Sequences:** WTD scaffold (UniProt Q863E9), all modeled variants (FASTA), and alignments (including ORdel mapping on the 24–233 axis).
- **RIDAO outputs:** per-residue disorder scores (24–233) for each sequence.
- **FuzDrop outputs:** per-residue p_DP_, S_bind_, protein-level p_LLPS_, and droplet-promoting region (DPR) segment lists.
- **KaML pK_a_ landscapes:** per-residue pK_a_ predictions (raw and WT-aligned Δ tables).
- **Discrete electrostatics:** integer-charge NCPR and charged-site counts at pH 7.0.
- **Figure source material:** input data along with scripts used to generate figures.
- **Structure inputs and visualization:** the 11 sequence-specific AlphaFold 3 models used as KaML-CBTree inputs, the WTD WT model shown in Fig. 2a, associated model-confidence outputs, and the corresponding PyMOL session file.

The MD trajectory-level numerical summaries, regional comparisons, interface occupancies, and contact-analysis definitions are provided with the Supplementary Information in the accompanying MD Supplementary Data files. The original simulation inputs, production trajectories, analysis scripts, and replicate-resolved numerical results are deposited separately on Zenodo and are publicly available at https://doi.org/10.5281/zenodo.23309128.

## CRediT authorship contribution statement

**GWD:** Conceptualization; Methodology; Software; Formal analysis; Investigation; Data curation; Visualization; Writing—original draft; Writing—review & editing. **LD**: Resources; Data curation; Validation; Writing—review & editing. **HW**: Resources; Interpretation; Writing—review & editing. **VNU:** Conceptualization; Methodology; Supervision; Interpretation; Writing—review & editing. **MS:** Conceptualization; Methodology; Supervision; Interpretation; Writing—review & editing; Funding acquisition; Project administration.

## Acknowledgments

This work was supported by Alberta Innovates, Grant 222300803.

