## Supplementary Information for "Predicted condensation-related signatures localize to two C-terminal regions of the prion protein across cervid PRNP variants associated with chronic wasting disease"

### Supplementary Methods

#### Additional sequence-level phase-separation predictors

The WTD reference and the 10 modeled variants analyzed in the main text were additionally evaluated using PScore, catGRANULES2, and DeePhase. PScore estimates phase-separation propensity from sequence-predicted planar $\pi$-contact frequencies.[1] catGRANULES2 and DeePhase provide sequence-level phase-separation scores using distinct sequence-derived feature sets.[2,3] Each predictor was run using its standard implementation and default settings.

For PScore, both the sequence-level score and residue-level profile were retained. For substitution variants, PScore sequence indices were mapped directly to the mature cervid PrP 24–233 coordinate frame. ORdel was mapped to the WT coordinate system using the same alignment procedure described in the main Methods; deleted WT positions 56–63 were treated as missing, and no values were interpolated. Residue-level PScore differences were calculated as variant minus WT at aligned positions.

PScore, catGRANULES2, and DeePhase provide additional sequence-level comparisons across the same variant panel. Because the predictors use different feature sets and output definitions, their scores were interpreted separately. The 126–140 and 220–230 focal intervals were selected from the residue-level FuzDrop regional summaries described in the main Methods.

#### Cross-species tyrosine correspondence

Tyrosine correspondences were used only to relate cervid positions to motifs reported in human PrP and did not affect any calculation. The following motif-level correspondences were used: WTD Y152/Y153 $\leftrightarrow$ human Y149/Y150 in helix $\alpha1$; WTD Y221 $\leftrightarrow$ human Y218 in the helix $\alpha3$ core; and WTD Y228/Y229 $\leftrightarrow$ human Y225/Y226 at the C-terminal end of helix $\alpha3$. No one-to-one human Tyr counterpart was assigned to WTD Y165/Y166; human Y169 was considered only as a nearby contextual residue.

Seven WTD Tyr residues—Y152, Y153, Y165, Y166, Y221, Y228, and Y229—were highlighted as literature-relevant structural reporters. Tyr remained neutral in the discrete-charge calculation, and predicted Tyr $pK_a$ shifts were interpreted qualitatively as additional structural context for the focal-region patterns.[4–8]

#### Atomistic molecular dynamics of monomeric PrP

**Construct and simulation design.** The independent molecular-dynamics (MD) panel comprised WT and the nine single-substitution variants (Q95H, G96S, A116G, M132L, S138N, S225F, S225Y, Q226K, and Q226E), each simulated in three independently initiated 600-ns trajectories. The monomeric construct contained cervid PrP residues 94–233; the disordered N-terminal residues 24–93 were not included, and the octarepeat deletion (ORdel) therefore was not simulated. The native Cys182–Cys217 disulfide was retained. Gly94 and Ala233 were modeled as neutral artificial termini. Histidine protonation assignments were fixed across systems (H99 HSE, H114 HSE, H143 HSD, H180 HSE, and H190 HSE). All systems originated from a common WT AlphaFold-derived starting backbone. Mutated side chains were introduced individually and the backbone N, C$\alpha$, C, and O coordinates restored to the WT starting values before topology generation to minimize mutation-independent initial differences.

GROMACS 2026.3[9] was used with the CHARMM36m protein force field,[10] CHARMM-modified TIP3P water, nominally 150 mM NaCl, 300 K, 1 bar, and a 2-fs production time step. Particle-mesh Ewald electrostatics and CHARMM force-switched Lennard–Jones interactions were used. A 10-ps position-restraint stage preceded production; production coordinates were written every 10 ps. Each independent trajectory spanned 0–600 ns, yielding 18 $\mu$s across 30 trajectories.

**Trajectory analyses.** Structural fitting used backbone atoms (N, C$\alpha$, C, O) in residues 144–153, 175–190, and 199–219, excluding both focal intervals. R1 (126–140) and R2 (220–

230) were specified from the sequence-based study before MD analysis. Measured descriptors included structured-core and whole-backbone RMSD, per-residue and regional backbone RMSF, residue solvent-accessible surface area (SASA), protein radius of gyration, DSSP secondary structure, and backbone $\phi/\psi$ distributions. Backbone distributions used $24 \times 24$ bins (15° in each dihedral) and base-2 Jensen–Shannon distance. RMSF was computed separately for each trajectory, rather than by concatenating coordinate trajectories. Protein-only analysis frames were sampled every 100 ps, giving 6,001 frames per trajectory including both endpoints.

Intramolecular hydrogen bonds used a 0.35-nm donor–acceptor distance cutoff and a 30° angle cutoff; donor–acceptor pairs differing only in hydrogen identity were merged. Salt bridges used a 0.40-nm minimum charged-atom distance between Asp/Glu side-chain oxygens and Lys/Arg side-chain nitrogens; neutral histidines were not included. The full R1×R2 interface comprised $15 \times 11 = 165$ predefined residue pairs, characterized using residue-pair-collapsed hydrogen-bond occupancy and all 165 C$\alpha$ distances.

A separate, exploratory heavy-atom physical-contact analysis counted a residue pair at most once per frame when any R1–R2 heavy-atom distance was less than or equal to the sum of the corresponding Bondi elemental van der Waals radii,[11] without added distance tolerance. The radii were 1.70, 1.55, 1.52, and 1.80 Å for carbon, nitrogen, oxygen, and sulfur, respectively. Periodic minimum-image distances were evaluated using MDAnalysis.[12] This additional measure characterizes intramolecular geometric contacts, not interaction free energies or independent confirmation of the hydrogen-bond measurements.

For intervals containing a substitution, the regional summaries were repeated with the mutated residue excluded; the same position was excluded from the WT comparator. Six consecutive 100-ns blocks were used as within-trajectory sampling diagnostics only. Equal-weight summaries across the three independent trajectories describe aggregate sampling (1.8 $\mu$s per system), while individual trajectories remain the replication units. The three WT trajectories define the observed WT sampling range; matching replicate numbers between WT and a variant are bookkeeping labels rather than a paired experimental design. No statistical tests treated frames or 100-ns blocks as independent replicates. These simulations model soluble monomeric PrP and do not model a condensed phase, intermolecular association, fibril formation, or CWD susceptibility.

### Supplementary Experimental Hypotheses and Context

#### Motif-centered tyrosine reporter hypotheses

**Helix $\alpha1$ nitration reporter.** Variants with higher predicted $p_{\mathrm{DP}}$ across residues 126–140 are hypothesized to alter the exposure or reactivity of the adjacent helix $\alpha1$ Tyr residues WTD Y152 and Y153, corresponding to human Y149 and Y150, during formation of $\beta$-rich aggregates. This hypothesis can be examined using targeted tyrosine nitration assays.[4,5]

**Helix $\alpha3$ nitration and shedding reporter.** Variants that lower $S_{\mathrm{bind}}$ within residues 220–230, including S225F and S225Y, are hypothesized to alter the reactivity of WTD Y228 and Y229, corresponding to human Y225 and Y226, during aggregation relative to WTD WT. Q226E is hypothesized to shift this pattern in the opposite direction. Because human Y226 is associated with the ADAM10 shedding site, this motif provides an additional biochemical reporter of local structural changes near the C-terminal end of helix $\alpha3$.[6,7]

#### PrP maturation and metal-dependent phase separation

Tange et al. showed that full-length recombinant PrP undergoes LLPS in a PEG/dextran aqueous two-phase system containing kosmotropic salts and that droplet aging can yield ThT-positive, sarkosyl-insoluble, proteinase K-resistant, $\beta$-sheet-rich gel-like material.[13] Deletion of the N-terminal region, residues 23–89, strongly reduced the increase in ThT signal during droplet maturation, and the aged recombinant PrP gel lacked RT-QuIC seeding activity. These findings distinguish LLPS and droplet maturation from the formation of templating-competent $PrP^{Sc}$.

Under the polymer–salt conditions used by Tange et al., $CuSO_4$ did not generate liquid droplets but instead produced amorphous aggregates and inhibited sulfate-induced droplet formation.[13] Metal-dependent condensation has been observed under other experimental conditions. Do Amaral et al. reported that $Cu^{2+}$ promoted PrP condensation in cells and in vitro, that preformed PrP:$Cu^{2+}$ condensates dissolved after EDTA chelation, that condensation was suppressed at pH 4.7, and that $H_2O_2$ promoted liquid-to-solid transition and amyloid-like aggregation.[14] A subsequent study found that $Zn^{2+}$ promoted recombinant PrP phase separation under defined near-neutral conditions and that this behavior depended on the N-terminal domain spanning residues 23–90.[15]

These studies show that metal effects on PrP phase behavior vary with buffer composition, pH, ionic strength, crowding, metal identity, and metal concentration. The near-neutral $pK_\mathrm{a}$ sites identified in residues 126–140 and 220–230 are therefore considered candidate modulators of local interaction modes, whereas $Cu^{2+}$ or $Zn^{2+}$ is not assumed to be generally required for PrP LLPS.

### Supplementary Tables

Table S1: Regional summaries across all non-WT variants

| **Region** | **Start** | **End** | med $\overline{\|\Delta p_{\mathrm{DP}}\|}$ | med $\overline{\|\Delta S_{\mathrm{bind}}\|}$ | med $\overline{\|\Delta pK_a\|}$ |
|---|---|---|---|---|---|
| 24–50 (CC1) | 24 | 50 | 0.0012 | 0.0846 | 0.014 |
| 51–91 (OR) | 51 | 91 | 0.0005 | 0.0622 | 0.011 |
| 92–120 (CC2–HR neighborhood) | 92 | 120 | 0.0167 | 0.1465 | 0.017 |
| 121–125 (pre-$\beta$1 linker) | 121 | 125 | 0.0248 | 0.1447 | – |
| 121–140 (pre-$\beta$1 linker + $\beta$1–$\alpha$1 region) | 121 | 140 | 0.0430 | 0.1771 | 0.015 |
| 126–140 ($\beta$1–$\alpha$1 region) | 126 | 140 | 0.0518 | 0.1409 | 0.015 |
| 141–143 ($\alpha$1–$\beta$2 linker) | 141 | 143 | 0.0231 | 0.1029 | 0.010 |
| 144–153 (helix $\alpha$1) | 144 | 153 | 0.0131 | 0.1224 | 0.022 |
| 154–161 (post-$\alpha$1 linker) | 154 | 161 | 0.0074 | 0.1096 | 0.009 |
| 162–171 ($\beta$2–$\alpha$2 loop) | 162 | 171 | 0.0098 | 0.0695 | 0.018 |
| 172–174 ($\beta$2–$\alpha$2 to $\alpha$2 N-cap linker) | 172 | 174 | 0.0170 | 0.0926 | 0.020 |
| 175–190 (helix $\alpha$2) | 175 | 190 | 0.0052 | 0.0420 | 0.021 |
| 191–198 ($\alpha$2–$\alpha$3 junction linker) | 191 | 198 | 0.0049 | 0.0157 | 0.010 |
| 199–206 ($\alpha$2–$\alpha$3 junction; $\alpha$3 N-cap) | 199 | 206 | 0.0044 | 0.0491 | 0.020 |
| 207–219 ($\alpha$3 hinge/linker) | 207 | 219 | 0.0057 | 0.1110 | 0.022 |
| 220–230 (helix $\alpha$3 C-terminal region) | 220 | 230 | 0.0291 | 0.1203 | 0.028 |
| 231–233 (C-terminal tail; pre-GPI signal) | 231 | 233 | 0.0295 | 0.2399 | 0.000 |

For each region and variant, the mean absolute WT-aligned difference was calculated across residues with valid values, and the resulting regional values were summarized using the median across the 10 non-WT sequences. For $pK_a$, only ionizable sites with valid KaML predictions were included. Rows are ordered by sequence position; no cross-metric composite score or overall rank was calculated. CC1, charged cluster 1; CC2, charged cluster 2; HR, hydrophobic region; OR, octarepeat region. The CC2–HR neighborhood denotes the 92–120 analytical interval, overlapping CC2 (95–111) and the N-terminal part of HR (112–133).

Table S2: Regional summaries for the protective subset

| **Region** | **Start** | **End** | med $\overline{\lvert\Delta p_{\mathrm{DP}}\rvert}$ | med $\overline{\lvert\Delta S_{\mathrm{bind}}\rvert}$ | med $\overline{\lvert\Delta pK_a\rvert}$ |
|---|---|---|---|---|---|
| 24–50 (CC1) | 24 | 50 | 0.0012 | 0.1334 | 0.014 |
| 51–91 (OR) | 51 | 91 | 0.0005 | 0.0889 | 0.010 |
| 92–120 (CC2–HR neighborhood) | 92 | 120 | 0.0167 | 0.1667 | 0.022 |
| 121–125 (pre-$\beta$1 linker) | 121 | 125 | 0.0285 | 0.1712 | – |
| 121–140 (pre-$\beta$1 linker + $\beta$1–$\alpha$1 region) | 121 | 140 | 0.0535 | 0.1847 | 0.015 |
| 126–140 ($\beta$1–$\alpha$1 region) | 126 | 140 | 0.0640 | 0.1628 | 0.015 |
| 141–143 ($\alpha$1–$\beta$2 linker) | 141 | 143 | 0.0194 | 0.1186 | 0.010 |
| 144–153 (helix $\alpha$1) | 144 | 153 | 0.0131 | 0.1329 | 0.022 |
| 154–161 (post-$\alpha$1 linker) | 154 | 161 | 0.0088 | 0.1213 | 0.009 |
| 162–171 ($\beta$2–$\alpha$2 loop) | 162 | 171 | 0.0098 | 0.0653 | 0.019 |
| 172–174 ($\beta$2–$\alpha$2 to $\alpha$2 N-cap linker) | 172 | 174 | 0.0186 | 0.0926 | 0.020 |
| 175–190 (helix $\alpha$2) | 175 | 190 | 0.0070 | 0.0420 | 0.021 |
| 191–198 ($\alpha$2–$\alpha$3 junction linker) | 191 | 198 | 0.0039 | 0.0077 | 0.010 |
| 199–206 ($\alpha$2–$\alpha$3 junction; $\alpha$3 N-cap) | 199 | 206 | 0.0032 | 0.0412 | 0.020 |
| 207–219 ($\alpha$3 hinge/linker) | 207 | 219 | 0.0061 | 0.1121 | 0.022 |
| 220–230 (helix $\alpha$3 C-terminal region) | 220 | 230 | 0.0344 | 0.1201 | 0.027 |
| 231–233 (C-terminal tail; pre-GPI signal) | 231 | 233 | 0.0227 | 0.2724 | 0.000 |

For each region and variant, the mean absolute WT-aligned difference was calculated across residues with valid values, and the resulting regional values were summarized using the median across the protective subset defined in the main text: Q95H, G96S, A116G, M132L, S138N, S225F, S225Y, and Q226K. This biological classification reflects published evidence of reduced susceptibility or case representation, delayed progression, or prolonged incubation, with different forms and strengths of evidence; it does not imply complete resistance. For $pK_a$, only ionizable sites with valid KaML predictions were included. Rows are ordered by sequence position; no cross-metric composite score or overall rank was calculated. CC1, charged cluster 1; CC2, charged cluster 2; HR, hydrophobic region; OR, octarepeat region. The CC2–HR neighborhood denotes the 92–120 analytical interval, overlapping CC2 (95–111) and the N-terminal part of HR (112–133).

Table S3: **Overlap summary: this work vs. Díaz-Domínguez *et al.*** Direction-only crosswalk for variants present in both this manuscript and Díaz-Domínguez *et al.*[16] For M132L, Díaz-Domínguez *et al.* report the substitution in a Q226E background, whereas this work models M132L on the WTD Q226 scaffold; arrows therefore provide qualitative context rather than a background-matched comparison for that row. Arrows indicate direction of change relative to the stated baseline (this work: WTD WT; Díaz: Q226Q reference, their Table 2).

| **Group** | **Variant** | **Díaz construct** | **This work (vs WT)** | | **Díaz (vs Q226Q)** | | **Díaz MD** |
|---|---|---|---|---|---|---|---|
| | | | $\Delta p_{\text{DP}}$ (126–140) | $\Delta S_{\text{bind}}$ (220–230) | $\Delta T_m$ | ΔScore | ΔRMSD |
| Ref | WT (WTD) | Q226Q | ∼ | ∼ | ∼ | ∼ | ∼ |
| 8V | Q95H (WTD) | Q95H Q226Q | ↑ | ∼ | ↑ | ↓ | NA[c] |
| 8V | G96S (WTD) | G96S Q226Q | ↑ | ∼ | ∼ | ↓ | NA[c] |
| 8V | A116G (WTD) | A116G Q226Q | ↑ | ↓[d] | ↑ | ∼ | NA[c] |
| 8V | M132L (elk context) | M132L Q226E | ∼ | ∼ | ↑ | ↓ | ↑ |
| 8V | S138N (reindeer/caribou) | S138N Q226Q | ↑ | ∼ | ↑ | ↓ | ↑ |
| 8V | S225F (mule deer) | S225F Q226Q | ↑ | ↓ | ↑ | ↓ | ↑ |
| 8V | S225Y (reindeer/caribou) | S225Y Q226Q | ↑ | ↓ | ↑ | ↓ | ↓ |
| 226E | Q226E (elk-typical) | Q226E Q226E | ∼ | ↑ | ↑ | ∼ | ↓ |
| 8V | Q226K (WTD) | Q226K Q226K | ↑ | ↓[d] | ↑ | ↓ | ∼ |

**Interpretive context for Table S3.** The comparison is qualitative because the two studies use different analytical readouts, reference baselines, and, for M132L, different residue-226 backgrounds. The global thermal-stability, molecular-dynamics, and spontaneous-misfolding measures reported by Díaz-Domínguez *et al.* do not reproduce the same variant ordering as the localized $p_{\text{DP}}$ and $S_{\text{bind}}$ signatures in this study. Q226E provides a clear example: it shows increased predicted $S_{\text{bind}}$ within residues 220–230 but no corresponding increase in the published spontaneous-misfolding score. This example shows that the localized predicted helix $\alpha 3$ $S_{\text{bind}}$ pattern and the global spontaneous-misfolding measure capture different aspects of variant behavior.

*Arrow key.* ↑ increase; ↓ decrease; ∼ no meaningful change.

*Baselines.* This work: relative to WTD WT on the 24–233 frame. Díaz-Domínguez *et al.*: relative to their reference construct Q226Q (their Table 2).

*Group key.* 8V = the protective subset defined in the main text by published evidence of reduced susceptibility or case representation, delayed progression, or prolonged incubation; the type and strength of evidence vary among variants and the label does not imply complete resistance. 226E = the elk-typical strain-context comparator Q226E; Ref = the WTD WT reference.

[c] NA indicates variants excluded from the molecular dynamics analysis of Díaz-Domínguez *et al.* because the polymorphisms fall exclusively within residues 23–131 and/or 230–234.

[d] Small-magnitude trend. "Small magnitude" denotes a negative trend near zero in this work's regional cross-metric summary; included for completeness.

*MD proxy.* ΔRMSD reflects the direction of change in the mean backbone RMSD reported by Díaz-Domínguez *et al.* for their $PrP^{C}$ molecular dynamics simulations.

Table S4: **Comparison of FuzDrop with three additional sequence-level phase-separation predictors**

| Sequence | FuzDrop $p_{LLPS}$ | PScore | ΔPScore vs WT | catGRANULES2 | ΔcatGRANULES2 vs WT | DeePhase | ΔDeePhase vs WT |
|---|---|---|---|---|---|---|---|
| WT | 0.7900 | 14.97 | 0.00 | 0.660 | 0.000 | 0.83 | 0.00 |
| Q95H | 0.7983 | 15.18 | +0.21 | 0.660 | 0.000 | 0.83 | 0.00 |
| G96S | 0.8768 | 14.78 | -0.19 | 0.685 | +0.025 | 0.84 | +0.01 |
| A116G | 0.9504 | 14.86 | -0.11 | 0.655 | -0.005 | 0.83 | 0.00 |
| M132L | 0.8340 | 14.91 | -0.06 | 0.680 | +0.020 | 0.84 | +0.01 |
| S138N | 0.8495 | 14.57 | -0.40 | 0.675 | +0.015 | 0.83 | 0.00 |
| S225F | 0.9012 | 14.97 | 0.00 | 0.680 | +0.020 | 0.83 | 0.00 |
| S225Y | 0.9072 | 14.97 | 0.00 | 0.700 | +0.040 | 0.82 | -0.01 |
| Q226K | 0.8125 | 14.97 | 0.00 | 0.675 | +0.015 | 0.83 | 0.00 |
| Q226E | 0.8513 | 14.97 | 0.00 | 0.655 | -0.005 | 0.84 | +0.01 |
| ORdel | 0.7159 | 11.97 | -3.00 | 0.690 | +0.030 | 0.82 | -0.01 |

FuzDrop $p_{LLPS}$, PScore, catGRANULES2, and DeePhase are predictor-specific sequence-level outputs with different score definitions. All Δ values are relative to WTD WT. The additional predictors provide method-specific comparisons across the sequence panel. The focal intervals were selected from the residue-level FuzDrop regional summaries described in the main Methods.

#### Comparison among sequence-level predictors

The additional predictors showed different sensitivities across the closely related PrP sequences (Table S4). PScore showed its largest change for ORdel, which decreased from 14.97 in WT to 11.97, whereas several substitutions at residues 225 and 226 produced no change in the sequence-level PScore. catGRANULES2 varied modestly across the panel, from 0.655 to 0.700, and DeePhase scores were narrowly distributed between 0.82 and 0.84. Thus, the three additional sequence-level predictors did not reproduce the variant ordering obtained from FuzDrop $p_{LLPS}$.

The different variant patterns are consistent with differences in predictor feature sets, training frameworks, score definitions, and sensitivity to individual substitutions. The sequence-level scores are therefore presented as method-specific comparisons. FuzDrop supplies the residue-resolved pDP and Sbind outputs analyzed in the main text, and the 126–140 and 220–230 focal intervals were selected from those regional summaries. Residue-level PScore profiles are shown in Fig. S4.

#### Regional and interregional MD results

Table S5: Regional backbone RMSF and intramolecular R1–R2 interface occupancy across all 30 trajectories

| Sequence | R1 RMSF (nm) | R2 RMSF (nm) | R1–R2 H-bond pairs/frame | R1–R2 physical contacts/frame |
|---|---|---|---|---|
| WT | 0.124 (0.109–0.149) | 0.148 (0.125–0.175) | 0.224 (0.130–0.318) | 0.384 (0.325–0.465) |
| Q95H | 0.150 (0.132–0.182) | 0.150 (0.142–0.162) | 0.555 (0.121–1.321) | 0.695 (0.252–1.436) |
| G96S | 0.179 (0.131–0.206) | 0.173 (0.145–0.227) | 0.286 (0.120–0.462) | 0.414 (0.260–0.587) |
| A116G | 0.142 (0.120–0.159) | 0.157 (0.145–0.179) | 0.457 (0.349–0.601) | 0.618 (0.549–0.734) |
| M132L | 0.117 (0.109–0.122) | 0.141 (0.129–0.155) | 0.263 (0.098–0.377) | 0.456 (0.293–0.546) |
| S138N | 0.130 (0.120–0.143) | 0.143 (0.133–0.162) | 0.300 (0.176–0.504) | 0.453 (0.331–0.641) |
| S225F | 0.175 (0.130–0.265) | 0.182 (0.153–0.215) | 0.835 (0.474–1.443) | 0.982 (0.620–1.581) |
| S225Y | 0.155 (0.134–0.172) | 0.167 (0.151–0.177) | 0.301 (0.110–0.576) | 0.466 (0.264–0.757) |
| Q226K | 0.134 (0.111–0.176) | 0.146 (0.130–0.170) | 0.143 (0.071–0.202) | 0.332 (0.295–0.386) |
| Q226E | 0.121 (0.104–0.143) | 0.144 (0.138–0.148) | 0.270 (0.203–0.362) | 0.413 (0.359–0.514) |

Each entry shows the mean of three independently initiated 600-ns trajectory means, followed by the observed range of those means. R1 = 126–140; R2 = 220–230. Each interregional residue pair is counted at most once per frame for the corresponding occupancy measurement; an H-bonded pair is defined separately from a heavy-atom physical contact.

In every trajectory, S225F and A116G showed larger R1–R2 heavy-atom physical-contact occupancy than the observed WT trajectory range. S225F also exceeded the WT range in interregional hydrogen-bond occupancy, including when the substituted residue 225 was excluded. These measurements reflect related aspects of an intramolecular interface, rather than two independent lines of evidence. Other variants showed greater overlap with WT or trajectory-dependent responses.

Table S6: Focal-region differences relative to WT in individual 600-ns trajectories

| **Variant** | **Region** | **ΔRMSF (nm)** | **ΔSASA (nm$^2$/res)** | **DSSP JS** | **$\phi/\psi$ JS** |
|---|---|---|---|---|---|
| | | *Each cell: replicate 1 / replicate 2 / replicate 3* | | | |
| Q95H | 126–140 | +0.027 / +0.033 / +0.017 | -0.007 / -0.004 / -0.015 | 0.129 / 0.243 / 0.139 | 0.193 / 0.292 / 0.168 |
| Q95H | 220–230 | +0.022 / -0.013 / -0.002 | +0.127 / -0.110 / -0.033 | 0.030 / 0.064 / 0.030 | 0.076 / 0.130 / 0.152 |
| G96S | 126–140 | +0.090 / +0.057 / +0.015 | +0.088 / +0.018 / +0.005 | 0.297 / 0.135 / 0.051 | 0.408 / 0.178 / 0.113 |
| G96S | 220–230 | +0.020 / +0.052 / +0.005 | -0.028 / -0.061 / +0.073 | 0.060 / 0.052 / 0.029 | 0.106 / 0.126 / 0.123 |
| A116G | 126–140 | +0.038 / +0.010 / +0.005 | +0.060 / -0.065 / -0.018 | 0.178 / 0.086 / 0.096 | 0.271 / 0.168 / 0.118 |
| A116G | 220–230 | +0.021 / +0.004 / +0.002 | +0.006 / -0.073 / -0.067 | 0.060 / 0.057 / 0.089 | 0.116 / 0.129 / 0.174 |
| M132L | 126–140 | +0.013 / -0.040 / +0.004 | +0.042 / -0.054 / -0.028 | 0.090 / 0.141 / 0.075 | 0.157 / 0.201 / 0.148 |
| M132L | 220–230 | +0.031 / -0.036 / -0.015 | -0.070 / +0.102 / -0.031 | 0.187 / 0.052 / 0.064 | 0.227 / 0.113 / 0.166 |
| S138N | 126–140 | +0.034 / -0.029 / +0.012 | +0.002 / -0.038 / +0.008 | 0.123 / 0.123 / 0.055 | 0.210 / 0.177 / 0.112 |
| S138N | 220–230 | +0.037 / -0.042 / -0.009 | +0.037 / +0.036 / -0.184 | 0.059 / 0.057 / 0.063 | 0.119 / 0.108 / 0.198 |
| S225F | 126–140 | +0.157 / -0.019 / +0.015 | +0.101 / -0.024 / -0.047 | 0.316 / 0.093 / 0.075 | 0.403 / 0.153 / 0.123 |
| S225F | 220–230 | +0.054 / +0.040 / +0.010 | -0.032 / +0.054 / +0.066 | 0.017 / 0.176 / 0.095 | 0.111 / 0.251 / 0.187 |
| S225Y | 126–140 | +0.051 / +0.023 / +0.018 | +0.019 / -0.018 / +0.002 | 0.244 / 0.088 / 0.118 | 0.323 / 0.164 / 0.209 |
| S225Y | 220–230 | +0.052 / -0.002 / +0.008 | +0.098 / +0.069 / -0.058 | 0.046 / 0.079 / 0.108 | 0.133 / 0.166 / 0.192 |
| Q226K | 126–140 | +0.067 / -0.038 / +0.000 | +0.056 / -0.064 / -0.083 | 0.129 / 0.144 / 0.081 | 0.227 / 0.184 / 0.125 |
| Q226K | 220–230 | +0.045 / -0.038 / -0.014 | +0.018 / +0.136 / +0.109 | 0.098 / 0.082 / 0.074 | 0.128 / 0.145 / 0.161 |
| Q226E | 126–140 | +0.034 / -0.034 / -0.011 | +0.092 / -0.085 / -0.095 | 0.124 / 0.131 / 0.104 | 0.204 / 0.185 / 0.116 |
| Q226E | 220–230 | +0.023 / -0.037 / +0.003 | -0.215 / +0.012 / +0.064 | 0.087 / 0.110 / 0.113 | 0.143 / 0.179 / 0.216 |

Signed values are variant minus WT, for each similarly labeled trajectory. Matching replicate labels are bookkeeping only, not evidence of a matched experimental design. DSSP and $\phi/\psi$ entries are regional Jensen–Shannon distances; their signs are not defined. These comparisons are descriptive and should be assessed alongside the three WT trajectories. Full sequence-wide and mutation-excluded measurements are provided in the accompanying MD numerical archive.

The sampled regional responses differ in magnitude and direction among independent trajectories. In particular, S138N R1 mobility differences overlap WT sampling variability and Q226E R2 solvent accessibility does not show a reproducible signed shift. A change in an MD descriptor is not taken as a direct measure of the FuzDrop droplet-promoting propensity or binding-mode entropy.

### Supplementary Figures

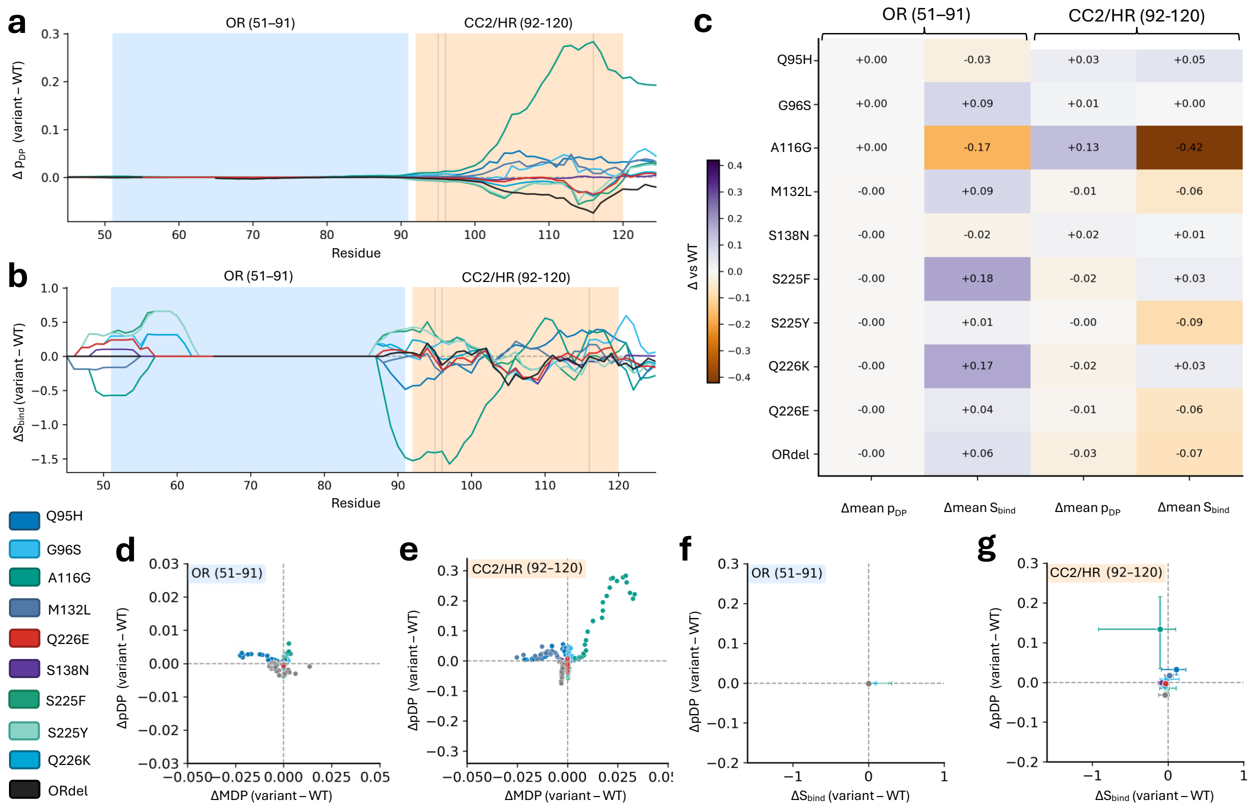


Figure S1: **N-terminal OR region and the defined CC2–HR neighborhood across variants.** (a) Per-residue change in droplet-promoting propensity, $\Delta p_{\mathrm{DP}}$ (variant–WT), across the octarepeat region (OR; residues 51–91) and the CC2–HR neighborhood (residues 92–120, overlapping charged cluster 2 and the hydrophobic region), highlighting that OR-region effects are comparatively small whereas CC2–HR effects can be variant-specific and larger in magnitude. (b) Per-residue change in binding-mode entropy, $\Delta S_{\mathrm{bind}}$ (variant–WT), across the same N-terminal segments, showing that the OR/CC2 transition exhibits the largest and most localized $\Delta S_{\mathrm{bind}}$ deviations among the N-terminal residues. (c) Heatmap summary of region-averaged $\Delta p_{\mathrm{DP}}$ and $\Delta S_{\mathrm{bind}}$ for the OR region (51–91) and the CC2–HR neighborhood (92–120), enabling direct comparison of region-level shifts across variants. (d,e) Parametric scatter of $\Delta p_{\mathrm{DP}}$ versus ΔMDP (variant–WT) for OR (d) and CC2–HR (e), indicating that OR points cluster near the origin while the CC2–HR neighborhood displays broader dispersion. (f,g) Parametric scatter of $\Delta p_{\mathrm{DP}}$ versus $\Delta S_{\mathrm{bind}}$ (variant–WT) for OR (f) and CC2–HR (g), showing that the larger N-terminal differences are concentrated after the OR and especially near the OR/CC2 transition. Colors denote variants and match the main-text palette; shaded backgrounds indicate the OR region (light blue) and the CC2–HR neighborhood (light orange).

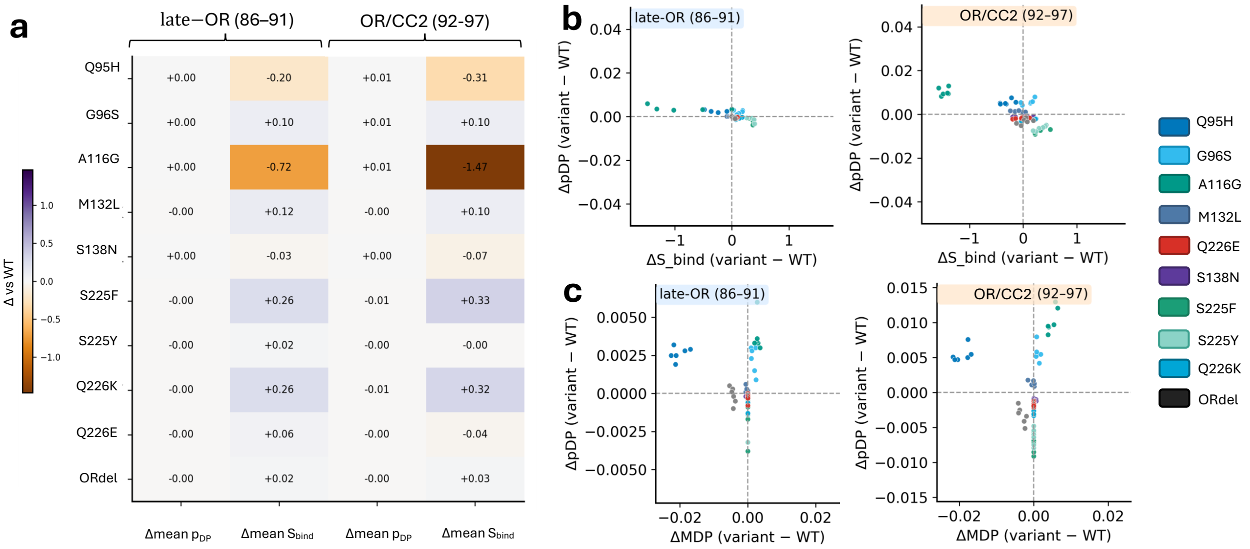


Figure S2: **Focused view of the OR/CC2 transition shows localized variant differences.** (a) Heatmap of region-averaged $\Delta p_{\mathrm{DP}}$ and $\Delta S_{\mathrm{bind}}$ for the C-terminal portion of the OR region (residues 86–91) and OR/CC2 transition (residues 92–97). Compared with averaging across the broader CC2–HR neighborhood (92–120; Fig. S1c), this focused analysis reduces dilution from flanking residues and emphasizes transition-local effects. (b) Parametric scatter of $\Delta p_{\mathrm{DP}}$ versus $\Delta S_{\mathrm{bind}}$ (variant–WT) for the C-terminal portion of the OR region (86–91; left) and OR/CC2 transition (92–97; right), illustrating greater variant-to-variant dispersion in the transition-adjacent segment than across the broader CC2–HR neighborhood. (c) Parametric scatter of $\Delta p_{\mathrm{DP}}$ versus $\Delta$MDP (variant–WT) for the C-terminal portion of the OR region (left) and OR/CC2 transition (right), providing an additional transition-local comparison consistent with the region-averaged heatmap in panel (a). Colors denote variants and match the main-text palette; region tags use the same OR (light blue) and CC2–HR (light orange) scheme.

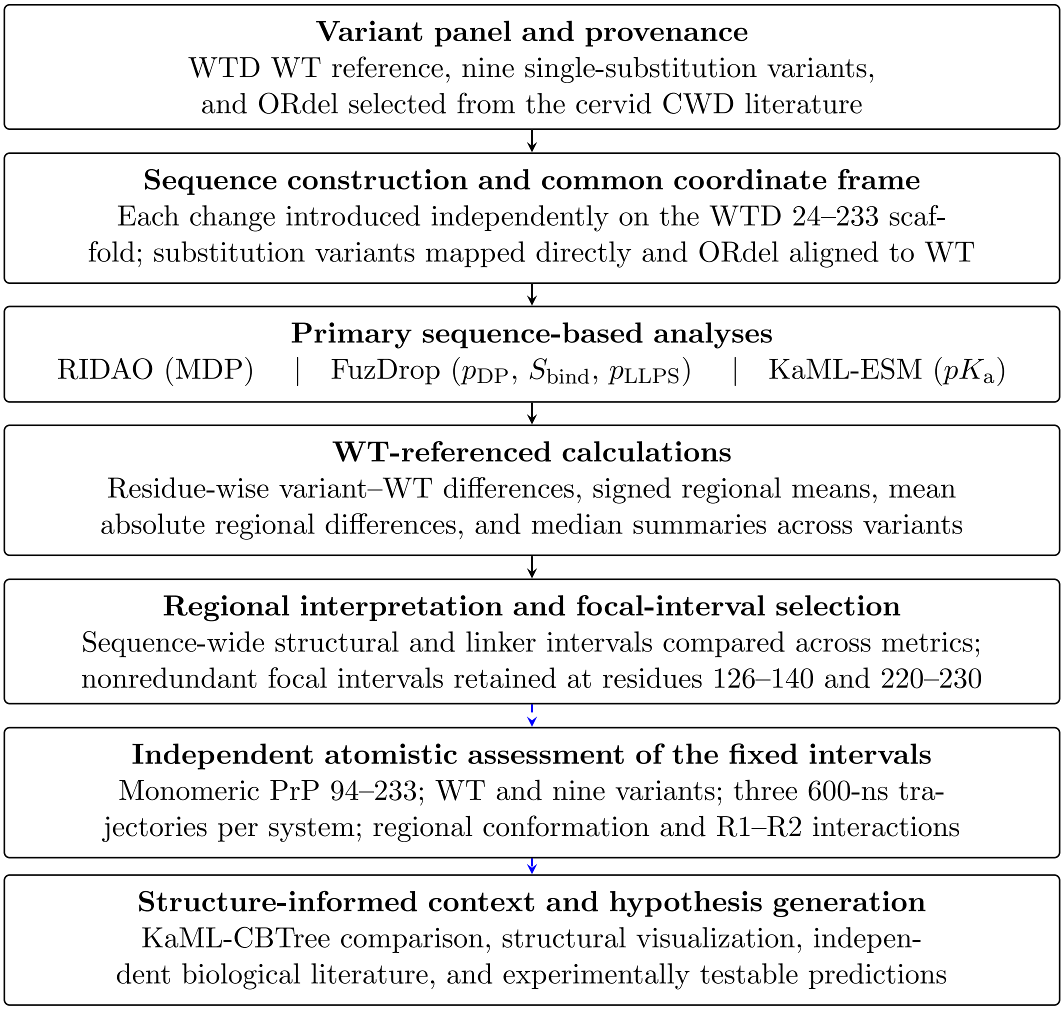


Figure S3: **Overview of the computational workflow.** Literature-supported cervid *PRNP* variants were modeled individually on a common WTD scaffold and analyzed using sequence-resolved disorder, condensation-related, and electrostatic predictors. Variant profiles were aligned to the WT coordinate system before residue-wise and regional differences were calculated. Sequence-wide regional summaries were used to prioritize nonredundant focal intervals for detailed interpretation. RIDAO, FuzDrop, and KaML-ESM generated the primary sequence-based outputs; KaML-CBTree, structural models, and published biological evidence provided context for interpretation and hypothesis generation. After the regions were fixed, independent atomistic MD examined monomeric conformational and interaction behavior; the simulations did not contribute to region selection.

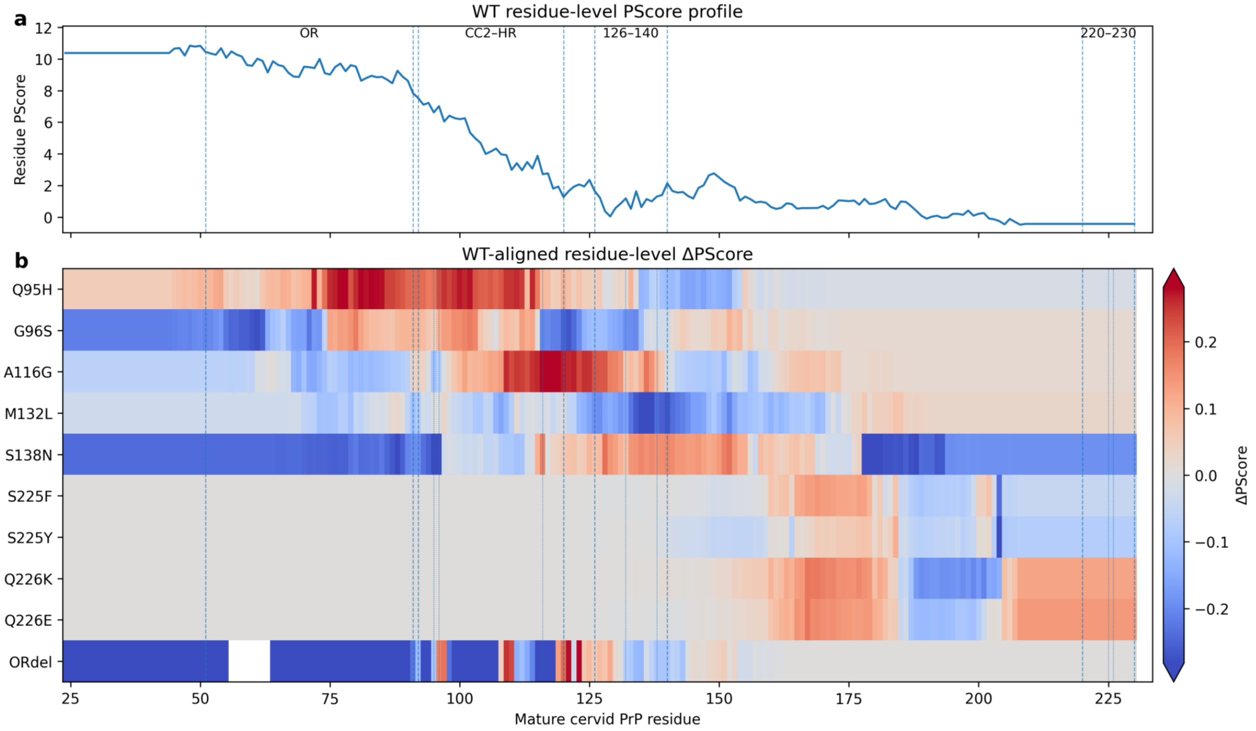


Figure S4: **Residue-level PScore profiles across cervid PrP variants.** **a**, WT residue-level PScore mapped to the mature cervid PrP 24–233 coordinate frame. Dashed boundaries indicate the octarepeat region, the CC2–HR neighborhood (92–120), and the focal intervals 126–140 and 220–230. **b**, WT-aligned residue-level PScore differences for the ten non-WT sequences. ORdel positions corresponding to deleted WT residues 56–63 are shown as missing. The diverging color scale was set using the single-substitution variants to preserve their contrast; ORdel values exceeding this range are saturated. PScore emphasizes the N-terminal sequence context and shows distributed responses to single substitutions, including localized C-terminal changes that may not alter the sequence-level PScore.

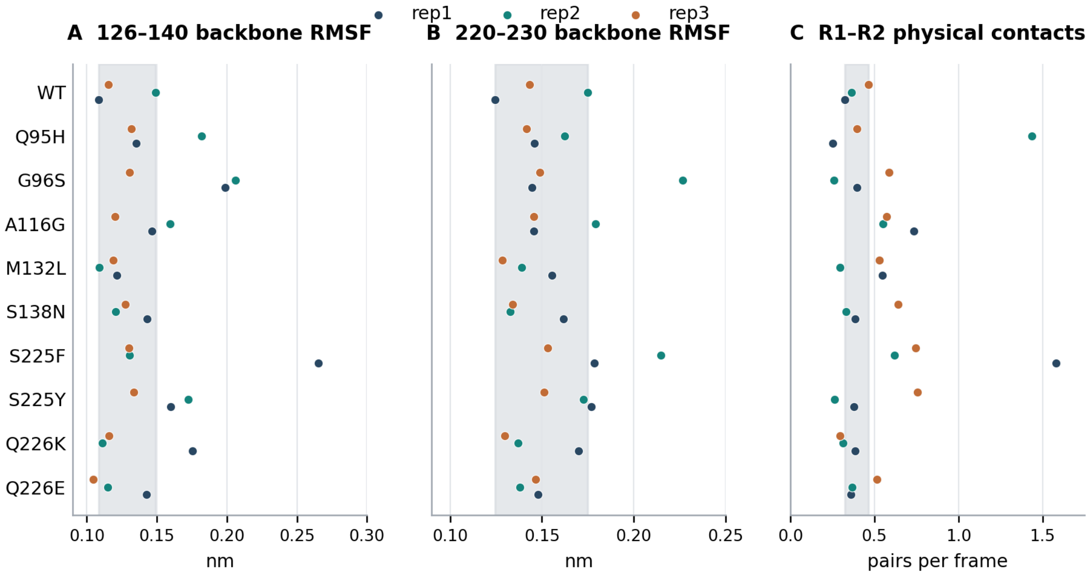


Figure S5: **Replicate-resolved MD comparisons across all nine substitutions and WT.** (**a,b**) Absolute backbone RMSF means in R1 (126–140) and R2 (220–230). (**c**) Mean number of contacting R1–R2 heavy-atom residue pairs per frame, counted using the sum-of-van-der-Waals-radii criterion. Each point represents one independent 600-ns trajectory; the three colors distinguish replicas. Gray bands span the observed WT trajectory-mean range for each descriptor. These structural observations describe monomeric PrP; frames and 100-ns blocks are not independent replicates.

**Supporting numerical data.** The companion MD Supplementary Data archive provides individual trajectory values plotted here, regional descriptor and mutation-excluded summaries, residue-pair interface occupancies, analysis definitions, and the heavy-atom contact-analysis script. The compact numerical archive does not contain the original production trajectories.

(15) do Amaral, M. J.; de Oliveira, L. S.; Cordeiro, Y. Zinc ions trigger the prion protein

liquid-liquid phase separation. *Biochemical and Biophysical Research Communications* **2025**, *753*, 151489.

(16) Díaz-Domínguez, C. M.; Eraña, H.; Peccati, F.; Vidal, E.; Charco, J. M.; Sampedro-Torres-Quevedo, C.; Pérez-Castro, M. A.; Lorenzo, N. L.; Giler, S.; Telling, G. C.; et al. Effect of primary structural variation on cervid prion protein in flexibility, stability, and spontaneous misfolding propensity. *Neurobiology of Disease* **2025**, *213*, 107005.
